# Hologenomic structure of bacterial and fungal community composition in the West Nile virus vector *Culex tarsalis*

**DOI:** 10.1101/2025.09.01.673577

**Authors:** Eunho Suh, Naomi Huntley, Travis van Warmerdam, Jamie P. Spychalla, Najmeh Nejat, Jason L. Rasgon

**Affiliations:** Department of Entomology, The Pennsylvania State University, University Park, PA, USA; Department of Biology, The Pennsylvania State University, University Park, PA, USA; One Health Microbiome Center, The Pennsylvania State University, University Park, PA, USA; Department of Plant Pathology and Environmental Microbiology, The Pennsylvania State University, University Park, PA, USA; Department of Biochemistry and Molecular Biology, The Pennsylvania State University, University Park, PA, USA; The Center for Infectious Disease Dynamics, The Pennsylvania State University, University Park, PA, USA; The Huck institutes of the Life Sciences, The Pennsylvania State University, University Park, PA, USA

**Keywords:** West Nile virus, microbiome, bacteria, fungi, mosquito, hologenome, RADSeq

## Abstract

**Background:** Microbiota play a crucial role in determining the ability for arthropod disease vectors to transmit pathogens. Microbial community structure can be heavily influenced by microbe-microbe interactions, host genetics and environmental factors. Here, we characterize the host population genetic structure, and bacterial and fungal communities in natural populations of the West Nile virus mosquito vector *Culex tarsalis*. Mosquitoes were collected and analyzed across the species range of the mosquito in the United States, where we used PoolRADSeq to quantify population genetic structure. Microbial community composition was characterized using bacterial 16S rRNA gene sequencing (V3-V4 region) and fungal ITS sequencing (ITS1 region).

**Results:** PoolRADSeq identified four broad genetic clusters of mosquito populations, which corresponded to previous clusters identified by microsatellite analysis and RADSeq on individual mosquitoes. Microbiome diversity grouped mosquito populations into three broad clusters, with each cluster distinctively represented by diagnostic abundant bacteria (*Ralstonia*, *Pseudomonas,* or *Zymobacter*/*Providencia*, respectively). Clustering for fungal taxa was less pronounced. Geographic distance between populations was positively correlated with microbiome community dissimilarity, and multiple environmental factors were significantly correlated with microbial species richness and diversity.

**Conclusions:** These results suggest that bacterial and fungal communities are geographically structured in *Cx*. *tarsalis*, interact with important environmental factors, and are partially correlated with host genetic structure. As microbiota can affect the ability for mosquitoes to transmit pathogens, understanding the factors underpinning microbiome variation across space and time has important implications for the spread of vector-borne pathogens such as WNV.

## Introduction

*Culex tarsalis* is one of the most important native North American mosquito arbovirus vectors [1]. The species is widely distributed in western North America, feeds primarily on avian hosts which contributes to virus amplification [2], and can move relatively long distances in search of a blood meal or oviposition sites [1,2,3]. Historically *Cx. tarsalis* is an important vector of Western equine encephalitis virus (WEEV) and Saint Louis encephalitis virus (SLEV) [4]. When West Nile virus (WNV) was introduced into the Americas in 1999, *Cx. tarsalis* was demonstrated to be one of the most competent vectors of this novel invasive pathogen [5].

Previously, population genetic data was used to implicate a role for *Cx. tarsalis* in transporting WNV across the western United States. An analysis of microsatellite loci suggested that across the western USA, *Cx. tarsalis* populations are structured into approximately three broad metapopulation clusters (“Midwest”, Sonoran [or “Southwest”], and “Pacific”), which closely matched the observed pattern of WNV geographic spread in the early 2000s [1,3]. A more recent population genetic analysis, using RADSeq on approximately 300 individual mosquitoes, identified a broadly similar pattern (with increased resolution and a new identified cluster [“Northwest”] due to the denser marker set) [6].

Knowledge of the factors influencing the spread of arboviruses in mosquito populations is critical for understanding dynamics of disease incidence, development of risk assessment strategies for novel virus introductions and development of virus transmission biomarkers that can be used to efficiently target control efforts [1,7]. In general, there are three main barriers that an arbovirus must overcome to be transmitted by a mosquito vector. The virus particles must 1) establish an infection of the midgut epithelium and replicate, 2) escape the midgut through the basal lamina and infect other body tissues, and 3) infect the salivary glands, escape into the salivary gland lumen and be transmitted during salivation [8,9,10]. These barriers can be modulated by the intrinsic host genetics of the mosquito, components of the endogenous mosquito microbiota, and environmental factors, which has been demonstrated for multiple mosquito species in both the lab and the field [11,12,13,14,15,16,17]. It has become increasingly clear that while pathogen transmission can be affected by host genetics and environmental factors, this is, at best, an incomplete representation of the factors influencing pathogen transmission phenotypes. Recent research on arthropod holobiomes (the host plus all associated microorganisms) has demonstrated the potential impact of mosquito-associated microorganisms on transmission-related phenotypes of a variety of pathogens [15,16,17,18,19,20,21,22, 23,24,25,26,27,28].

Laboratory studies have demonstrated that components of the mosquito holobiome such as bacteria and fungi can modulate pathogen vector competence phenotypes. For example, bacteria in the genera *Chromobacterium*, *Enterobacter*, *Serratia*, *Pseudomonas,* and *Delftia* can decrease *Plasmodium* oocyst levels in *Anopheles* mosquitoes [17,20,21,23], while bacteria in the genera *Serratia* and *Aeromonas* can increase mosquito susceptibility to arboviruses [22,24]. Less is known about how fungi alter pathogen vector competence in mosquitoes, but examples exist. Specific isolates of *Penicillium* enhanced *Anopheles* infection with *Plasmodium* [25], while *Metarhizium anisopliae* infection rendered *Aedes* mosquitoes less susceptible to dengue virus [26]. These data suggest that components of the mosquito microbiome can be potent modulators of pathogen transmission, but despite their importance there is very little data on what role these microorganisms play in determining vector competence variation in the field.

Little is known about the natural microbiome of *Cx*. *tarsalis* or its effects on pathogen infection and transmission. Current studies are limited to characterizing bacterial communities in *Cx. tarsalis* mosquitoes reared in experimental mesocosms, where communities were dominated by the aquatic bacteria *Thorsellia* [29, 30]; a common microbe found in freshwater aquatic environments [30]. Nothing is currently known about fungal communities in this species. To address this knowledge gap, we examined the bacterial and fungal communities in wild-caught *Cx. tarsalis* mosquitoes from populations across the species range in the western USA. Combined with host RADSeq data, we found signatures of population structuring in microbial communities that were partially correlated with mosquito population genetic structure, where mosquitoes clustered into three broad groups based on their microbial composition. Microbial structure was more pronounced for bacteria than fungi, possibly reflecting differences in microbe acquisition mechanisms. Geographic distance between populations was positively correlated with microbial community dissimilarity. Environmental factors such as longitude, latitude, temperature, and precipitation were significantly correlated with species richness and/or diversity. These results suggest that bacterial and fungal communities in this mosquito are geographically structured, are potentially shaped by host genetics, and interact with or are affected by environmental factors. These results have critical significance for understanding patterns of microbial ecology in this mosquito species, as well as having implications for the epidemiology of vector-borne pathogens such as West Nile virus.

## Methods

### PoolRADSeq

2b-RADseq on pooled mosquitoes was used to characterize *Cx. tarsalis* population structure. Wild *Cx. tarsalis* mosquitoes were collected from 15 populations in 2021 and 18 populations in 2022 using CDC traps (Fig 1, Table 1). Mosquitoes were shipped overnight on dry ice to Penn State and stored at −80 °C until processing. For each population, we created pools of 10 randomly selected mosquitoes. DNA was extracted from individuals using Qiagen spin columns, quantified with Qubit, and equal amounts of DNA from each individual were combined to form each pool. After DNA QC, standard 5’ –NNN-3’ adaptors were used to ligate restriction tags for concatenated Isolength restriction site-associated tags sequencing. Genomic DNA was digested with the type IIB restriction enzyme BsaXI to produce restriction fragments of uniform length, followed by adaptor ligation, amplification, purification, concatenating and barcoding to incorporate sample-specific barcodes. Samples were then pooled, QC assessed, and sequenced on the Illumina Hiseq Xten/NovaSeq platform using paired end 150bp sequencing. Reads were quality filtered. Reads containing the restriction endonuclease recognition site were extracted from the sequencing data and clustering was performed with ustacks software (version 1.34) from the Stacks package to construct a reference sequence. Sequencing data was aligned to the reference sequence using SOAP software (version 2.21), and SNP calling was performed using Maximum Likelihood (ML) using the RADtyping package [31]. A total of 9,626 quality-filtered SNP markers were obtained across all samples. Phylogenetic trees were constructed using Neighbor Joining. Library construction, sequencing, and analysis were conducted by Novogene.

**Fig. 1.**
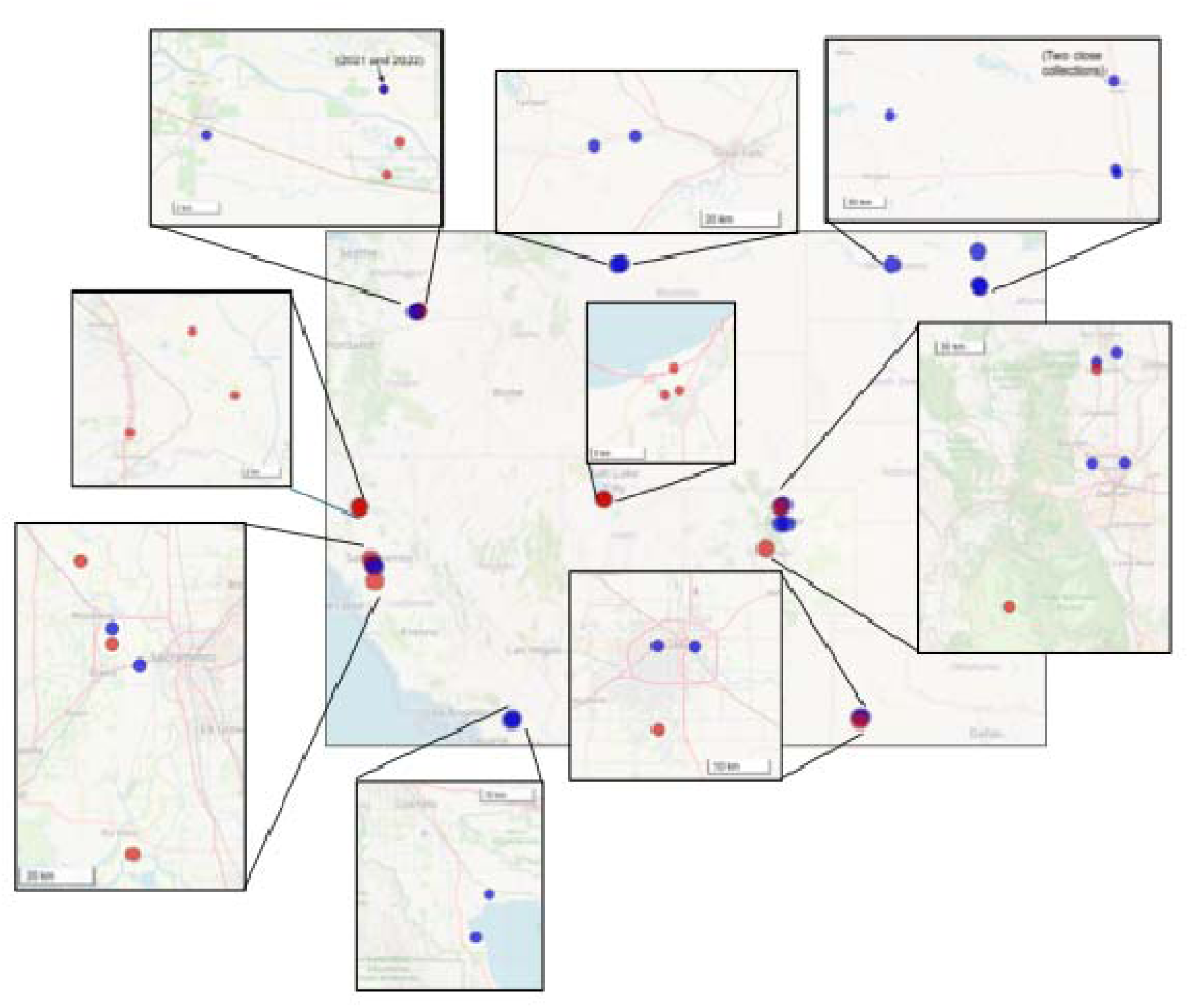
Map of collection locations for mosquitoes assayed for PoolRADSeq analysis. Red = 2021; Blue = 2022.

**Table 1.** Collection information for PoolRADSeq samples.

| Collection year | Sample ID | Location | State | Latitude | Longitude |
| --- | --- | --- | --- | --- | --- |
| 2021 | P1_2021 | Lake point | UT | 40.669406 | -112.305263 |
| 2021 | P2_2021 | Stansbury Park | UT | 40.651018 | -112.299081 |
| 2021 | P3_2021 | Stansbury Park | UT | 40.647411 | -112.314970 |
| 2021 | P4_2021 | Yakima/Grandview | WA | 46.193421 | -119.899400 |
| 2021 | P5_2021 | Yakima/Grandview | WA | 46.224977 | -119.901053 |
| 2021 | P6_2021 | Yakima/Grandview | WA | 46.205611 | -119.892026 |
| 2021 | P7_2021 | Lubbock | TX | 33.462700 | -101.891400 |
| 2021 | P9_2021 | Loveland | CO | 40.403933 | -105.071421 |
| 2021 | P10_2021 | Loveland | CO | 40.401394 | -105.108170 |
| 2021 | P12_2021 | Cottonwood, Shasta County | CA | 40.414286 | -122.216945 |
| 2021 | P13_2021 | Anderson, Shasta County | CA | 40.445232 | -122.243092 |
| 2021 | P14_2021 | Cottonwood, Shasta County | CA | 40.396727 | -122.281994 |
| 2021 | P15_2021 | Yolo County | CA | 38.610336 | -121.711942 |
| 2021 | P16_2021 | Sacramento County | CA | 38.107311 | -121.649758 |
| 2021 | P17_2021 | Yolo County | CA | 38.810503 | -121.808285 |
| 2022 | Z1 | Riverside county, Mecca | CA | 33.53271 | -116.03503 |
| 2022 | Z2 | Riverside county, Thermal | CA | 33.46238 | -116.06125 |
| 2022 | Z3 | Yolo county, Conaway Ranch | CA | 38.64849 | -121.71102 |
| 2022 | Z4 | Yolo county, Vic Fazio | CA | 38.56179 | -121.62924 |
| 2022 | Z5 | Adams County | CO | 39.90015 | -104.91565 |
| 2022 | Z6 | Broomfield | CO | 39.89499 | -105.13878 |
| 2022 | Z7 | Larimer county, Loveland | CO | 40.43432 | -105.10429 |
| 2022 | Z8 | Larimer county, Fort Collins | CO | 40.48614 | -104.97362 |
| 2022 | Z9 | Mapleton | ND | 46.89102 | -97.04281 |
| 2022 | Z10 | Mapleton | ND | 46.88508 | -97.06925 |
| 2022 | Z11 | Grand Forks county, Grand Forks | ND | 46.94475 | -97.06344 |
| 2022 | Z12 | Grand Forks county, Grand Forks | ND | 47.88686 | -97.09869 |
| 2022 | Z13 | Cascade county, Sunriver | MT | 47.51992 | -111.77331 |
| 2022 | Z14 | Cascade, Vaughan | MT | 47.53991 | -111.63364 |
| 2022 | Z15 | Yakima county, Grandview | WA | 46.22498 | -119.90105 |
| 2022 | Z16 | Yakima county, Mabton | WA | 46.20787 | -119.99583 |
| 2022 | Z17 | Lubbock, Lubbock | TX | 33.58318 | -101.89314 |
| 2022 | Z18 | Lubbock, Lubbock | TX | 33.58254 | -101.82794 |

### Microbiome mosquito collection and DNA extraction

*Cx. tarsalis* mosquitoes were collected from 17 locations in five states (CA, CO, UT, TX, and WA) in 2021 (Fig 2, Table 2). Mosquitoes were collected using CO_2_ baited EVS (encephalitis vector surveillance) or CDC traps overnight and morphologically identified at species level by the collecting laboratory. Mosquito samples were stored in DNA/RNA Shield (Zymo Research, Irvine, USA) at −80C until they were shipped to Penn State University. Morphological species identification of *Cx. tarsalis* was confirmed, and individual mosquitoes were surface sterilized in 500 µl 70% ethanol for 5 minutes followed by 3 washes in PBS. All tools used for handling mosquitoes were sterilized using absolute ethanol to minimize cross contaminations. DNA extractions were performed using ZymoBiomics DNA/RNA miniprep kit (Zymo Research, Irvine, USA) following the manufacture’s protocol with some modifications. In brief, individual whole mosquitoes were homogenized in 2.0mm Zymo BeadBeating Lysis tubes (Zymo Research, Irvine, USA) for 30 second at 6m/s in 750ml Zymo DNA/RNA Shield lysis buffer (Zymo Research, Irvine, USA) using a Fisher brand Beadmill 24 (Fisher Scientific, Waltham, MA) followed by additional mechanical homogenization in 0.1-0.5mm BeadBeating Lysis tubes for two 30 second cycles at 6m/s with a 45 second dwell. The homogenization protocol was optimized to ensure extraction of both bacterial and fungal DNA from a single mosquito while minimizing DNA degradation by excessive shearing. To minimize batch effects, two randomly selected populations were processed per day by two personnel, with a blank (PBS) extraction added as negative control for each batch (total 16 negative controls). All negative controls were determined to be negative for 16S PCR and did not pass QC for library preparation. For positive control, two units of ZymoBIOMICS Microbial Community Standard (Zymo Research, Irvine, USA) were used to validate our extraction method by confirming detection of known bacterial and fungal species in the standard mixture. Quantity and quality of DNA were checked using Qubit 4.0 fluorometer (Fisher Scientific, Waltham, USA) and nanodrop, respectively. DNA samples were stored at −80C until down-stream processing. Sample sizes and collection locations are listed in Table 2.

**Fig. 2.**
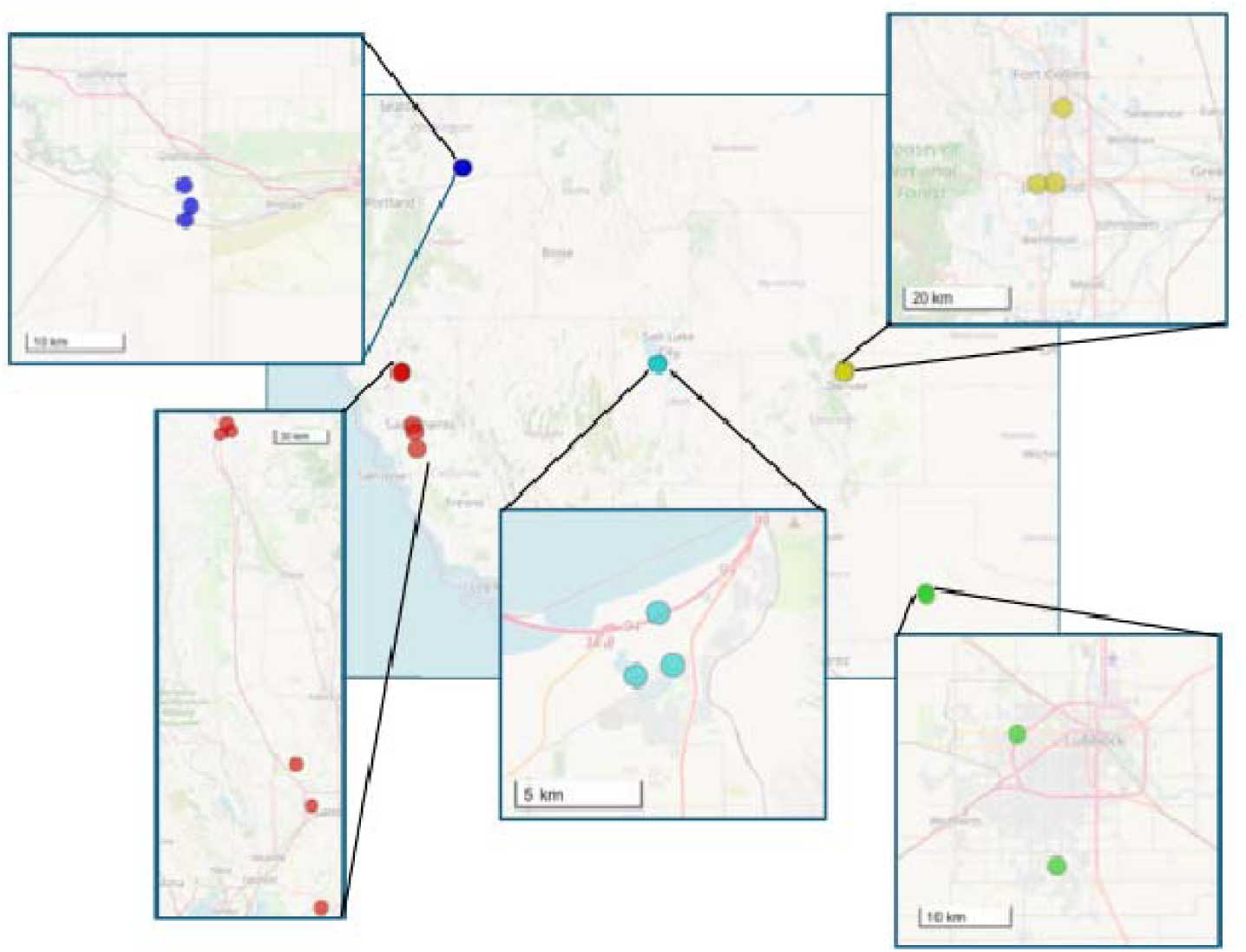
Map of collection locations for mosquitoes assayed for microbiome analysis. Colors indicate States.

**Table 2.**
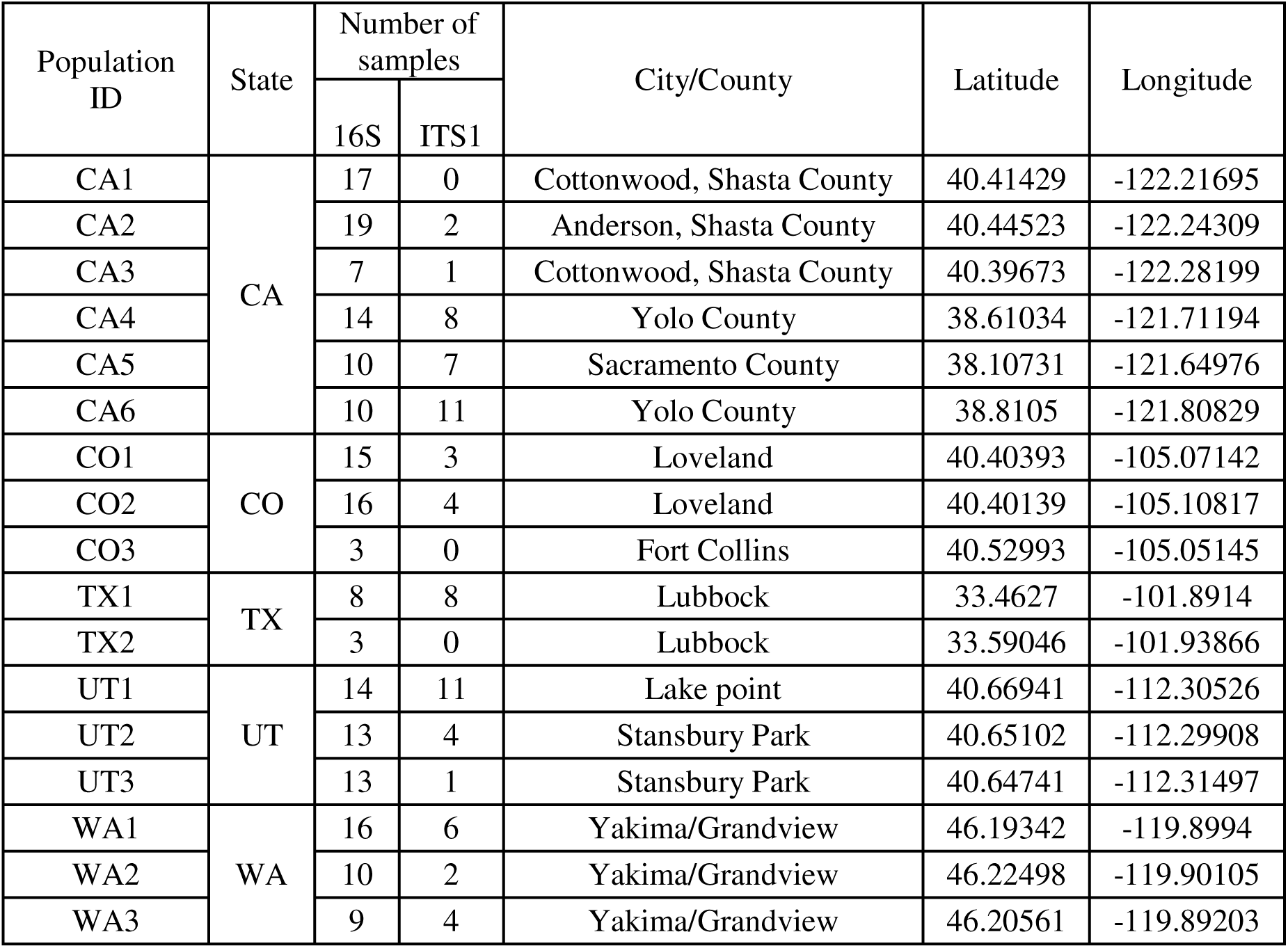
Collection information for microbiome samples.

| Population ID | State | Number of samples |  | City/County | Latitude | Longitude |
| --- | --- | --- | --- | --- | --- | --- |
|  |  | 16S | ITS1 |  |  |  |
| CA1 | CA | 17 | 0 | Cottonwood, Shasta County | 40.41429 | -122.21695 |
| CA2 |  | 19 | 2 | Anderson, Shasta County | 40.44523 | -122.24309 |
| CA3 |  | 7 | 1 | Cottonwood, Shasta County | 40.39673 | -122.28199 |
| CA4 |  | 14 | 8 | Yolo County | 38.61034 | -121.71194 |
| CA5 |  | 10 | 7 | Sacramento County | 38.10731 | -121.64976 |
| CA6 |  | 10 | 11 | Yolo County | 38.8105 | -121.80829 |
| CO1 | CO | 15 | 3 | Loveland | 40.40393 | -105.07142 |
| CO2 |  | 16 | 4 | Loveland | 40.40139 | -105.10817 |
| CO3 |  | 3 | 0 | Fort Collins | 40.52993 | -105.05145 |
| TX1 | TX | 8 | 8 | Lubbock | 33.4627 | -101.8914 |
| TX2 |  | 3 | 0 | Lubbock | 33.59046 | -101.93866 |
| UT1 | UT | 14 | 11 | Lake point | 40.66941 | -112.30526 |
| UT2 |  | 13 | 4 | Stansbury Park | 40.65102 | -112.29908 |
| UT3 |  | 13 | 1 | Stansbury Park | 40.64741 | -112.31497 |
| WA1 | WA | 16 | 6 | Yakima/Grandview | 46.19342 | -119.8994 |
| WA2 |  | 10 | 2 | Yakima/Grandview | 46.22498 | -119.90105 |
| WA3 |  | 9 | 4 | Yakima/Grandview | 46.20561 | -119.89203 |

### Microbiome sequencing and bioinformatics analysis

Bacterial and fungal communities were characterized by splitting the DNA from same individual mosquito. For bacterial characterization, the V3-V4 hypervariable region of 16S rRNA gene was amplified using primers 341F (5’-CCTAYGGGRBGCASCAG-3’) and 806R (5’-GGACTACNNGGGTATCTAAT-3’) with expected amplicon size of approximately 450-550bp. For fungal characterization, the Internal Transcribed Spacer (ITS) 1 region was amplified using primers ITS1F (5’-CTTGGTCATTTAGAGGAAGTAA-3’) and ITS2 (5’-GCTGCGTTCTTCATCGATGC-3’) with expected amplicon size of approximately 200-400bp. PCR was carried out in 30 μL reactions with 15 μL of Phusion High-Fidelity PCR Master Mix (New England Biolabs), 0.6 mM of forward and reverse primers, and 5∼10 ng of template DNA (10 μL for negative controls). Thermal cycling for bacterial characterization consisted of initial denaturation at 98°C for 1 min, followed by 30 cycles of denaturation at 98°C for 10 sec, annealing at 50°C for 30 sec, and extension at 72°C for 30 sec, and lastly by 72°C for 5 min. The same cycle was used for fungal characterization, except for the annealing temperature of 55°C for 30 sec. PCR products within the expected size were selected by 2% agarose gel electrophoresis. The same amount of PCR product from each sample was pooled, end-repaired, A-tailed and further ligated with Illumina adapters. PCR products were purified using the Qiagen Gel Extraction Kit (Qiagen, Germany). Sequencing libraries were generated using NEBNext Ultra DNA Library Pre ®Kit for Illumina. Libraries were commercially sequenced (Novogene) to generate 250 bp paired-end raw reads. The library was checked by Qubit and real-time PCR for quantification, and bioanalyzer for size distribution detection. Quantified libraries were pooled and sequenced on the NovaSeq PE250 platform (Illumina, San Diego, CA, USA).For sequencing and data processing, paired-end reads were assigned to samples and truncated by cutting off the barcode and primer sequences. After truncation, FLASH (v1.2.11, ccb.jhu.edu/software/FLASH/) was used to merge reads to get raw tags [2]. Then, fastp software was used to perform quality control of raw tags, and high-quality clean tags were obtained. Finally, chimeras were removed using VSEARCH. For denoising and species annotation of ASVs (Amplicon Sequencing Variants), DADA2 was used to reduce noise and sequences with abundance less than 5 were filtered out to obtain the final ASV [32]. Representative sequences for each ASV were annotated to obtain the corresponding species identification using QIIME2. By applying QIIME2’s classify-sklearn algorithm [33,34], a pre-trained Naive Bayes classifier was used for species annotation of each ASV. PCR, library preparation, sequencing and data processing were performed by Novogene.

### Microbiome data and statistical analysis

Data and statistical analyses were conducted using MicrobiomeAnalyst 2.0 [35], GraphPad Prism (v8.0), and R. For data filtering prior to downstream analyses, features that were constant across all samples and singletons (those with only one total count) were removed. To eliminate low-quality or uninformative features, the low-count filter was set to zero to avoid random selection bias for tied low-count data, and the low-variance filter was set to 10% interquartile range, followed by a data normalization process. Filtered data were used for all downstream comparative analyses unless otherwise specified. To understand overall bacterial and fungal community structure, we first constructed relative abundance and heat trees of representative taxa. Based on visual inspection of these data, we hypothesized a clustering of mosquito populations and validated this grouping through a series of comparative analyses including Alpha and Beta diversity, Random Forest classification, Core microbiome analysis, dendrogram construction, Linear Discriminant Analysis Effect Size (LEfSe), correlation network analysis, and correlations between environmental factors, community structure, and geographic distance. For Alpha diversity analysis, non-filtered original data were used. For Random Forest classification, five consecutive Out-of-Bag (OOB) error tables were generated to estimate the mean OOB error. Correlation network analysis employed Sparse Estimation of Correlations among Microbiomes (SECOM) with Pearson1, which efficiently handles both linear and non-linear relationships among microbes while maintaining sparsity. Correlation analyses between geographical distance and community dissimilarity indices were performed using the Mantel test (*vegan* package) in R. All other analyses performed in MicrobiomeAnalyst 2.0 followed the default settings as recommended [35]

### *Wolbachia* verification

*Wolbachia* was detected by 16S amplicon sequencing in a few (12/197) samples. For these samples we attempted to verify this finding by amplifying and sequencing the *Wolbachia* surface protein (*wsp*) gene using primers 81F and 691R [36]. We also amplified and sequenced the mosquito mitochondrial ND4 gene [37] to verify mosquito species identification. PCR and Sanger sequencing for these genes was carried out as described in the respective references.

## Results

### RADSeq population genetics

A total of 9,626 SNP markers that passed quality control were obtained across all samples, which were distributed across UTRs, introns, exons, and intergenic regions. Results from both years were combined for analysis to obtain a more complete spatial sampling across the species range of the mosquito. Clustering neighbor-joining results from PoolRADSeq were qualitatively similar to previous results from RADSeq on individual mosquitoes [6] and microsatellites [1,3], resolving the same metapopulation clusters (Pacific, Sonoran/Southwest, Northwest, and Midwest) that previous analyses [1,3,6] identified (Fig 3).

**Fig. 3.**
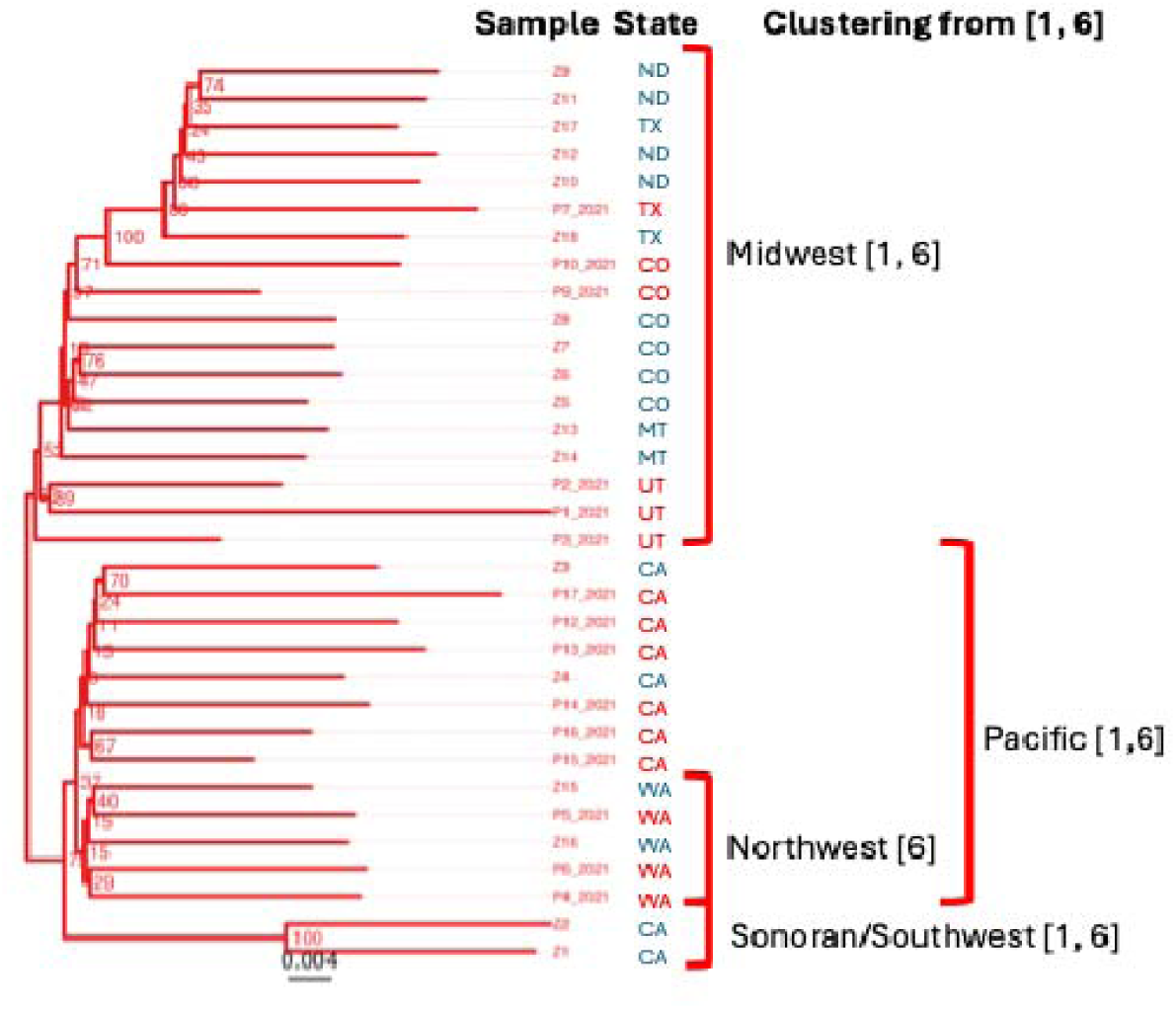
Neighbor-joining tree generated from PoolRADSeq analysis. Clusters previously identified by Venkatesan and colleagues (microsatellites) [1] and Liao and colleagues (individual RADSeq) [6] are highlighted.

### Microbiome composition profile

For processing bacterial sequence data, total read count was 27,771,776, and the average count per sample was 140,973 (Maximum = 502,316; Minimum = 10,132) from 197 samples that passed QC. A total 107 ASVs were left of 66396 by removing singletons (one total count) and low abundance ASVs (less than 4 read count). For fungal sequence data, total read count was 16,402,771 and the average count per sample was 227,816 (Maximum = 792,580; Minimum = 23,624) from 72 samples that passed QC. A total of 42 ASVs were left of 8,261 by removing singletons and low abundance ASVs. For bacterial and fungal combined sequence data, total read count was 26,776,978 and the average count per sample was 377,140 (Maximum = 1,001,576; Minimum = 76,642) from 71 samples. 163 ASVs were left of 74,657 by removing singletons and low abundance ASVs.

We first investigated overall community composition in bacteria and fungi in across all *Cx*. *tarsalis* populations. For bacterial communities, *Proteobacteria* were the most abundant at the phylum level (89.8%) which was mainly composed of *Gammaproteobacteria* (85.7% of total abundance) (Fig 4; Additional file 1). Within *Gammaproteobacteria*, *Burkhoderiales* was most abundant followed by *Pseudomonadales*, *Enterobacterales* and *Oceanospirillales*, representing over 85% of total abundance at the order level. Within *Gammaproteaobacteria*, *Ralstonia*, *Pseudomonas*, *Zymobacter*, and *Providencia* were the most abundant taxa representing 75% of total abundance at the genus level. While the top three abundant taxa were found in 74% to 96% of all 197 samples, *Providencia* was relatively less prevalent (25%).

**Fig. 4.**
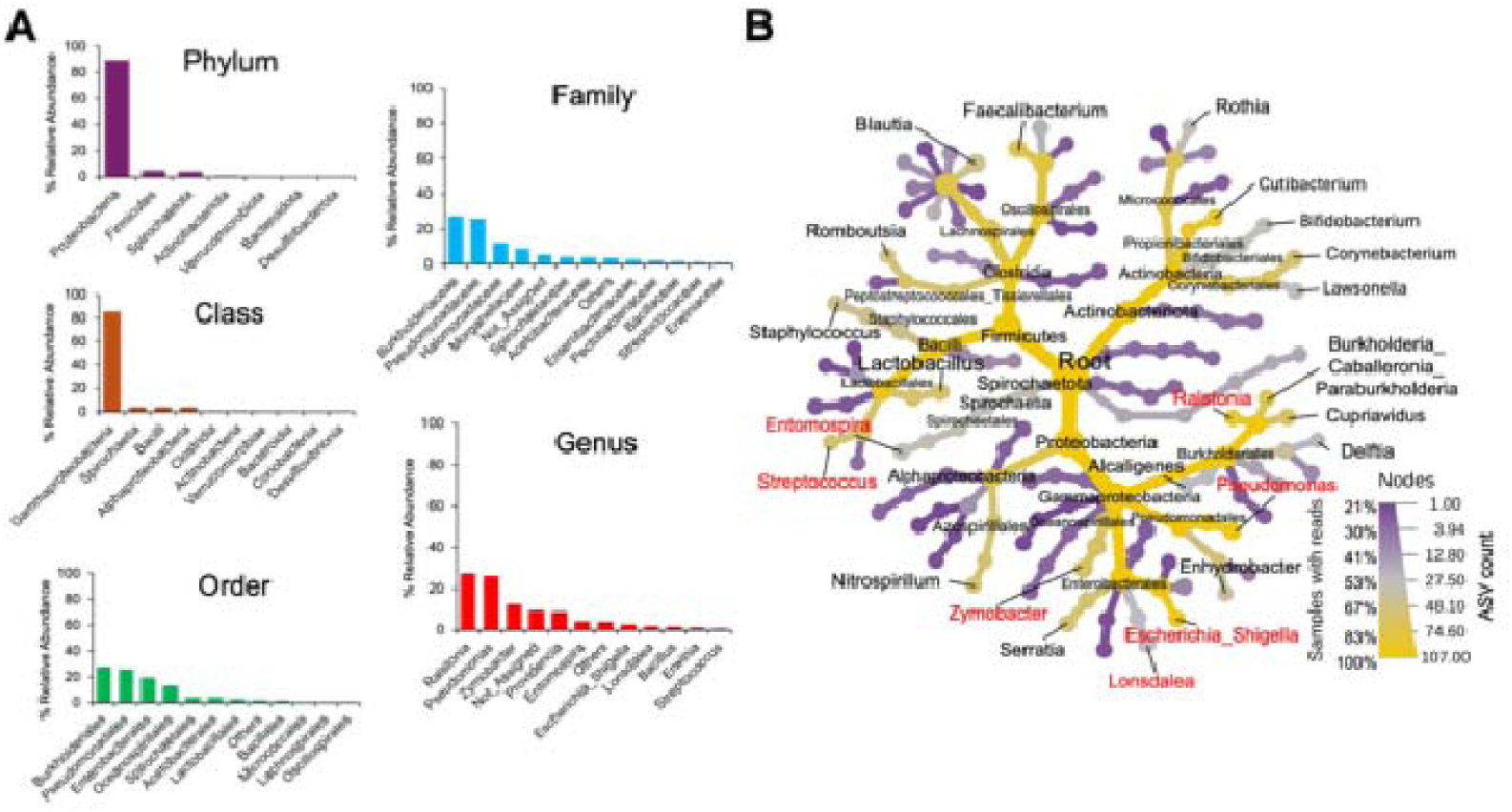
Bacterial communities in *Culex tarsalis* mosquitoes. **A:** Percent relative abundance of the top 10 identified taxa. **B:** Heat tree showing hierarchical structure of representative taxa. Taxa with 50% prevalence (% ASV detected in all samples) or higher are presented (family names are not labeled for better visibility). Genera within the top 10 relative abundance (see Fig. 1A) are in red text. Color and node indicate prevalence and number of ASV, respectively. See Additional file 1 for heat tree with all taxa represented.

We detected *Wolbachia* in approximately 6% (12/197) of samples. In previous surveys, *Cx. tarsalis* was found to be uninfected with *Wolbachia* [39], so we attempted to further validate this finding. From these 12 samples, we amplified and sequenced the *Wolbachia wsp* gene to strain type the infection. PCR was successful in only 2 samples, where the infection was identified as *w*Pip (the canonical *Wolbachia* strain infecting *Cx. pipiens/quinquefasciatus*; [40]); the remaining 10 samples either did not amplify or amplified random mosquito DNA. We then amplified and sequenced the mosquito mitochondrial ND4 gene to molecularly identify the mosquitoes. The two verified *Wolbachia*-positive samples were identified as *Cx*. *quinquefasciatus*. The other 10 samples were identified as *Cx. tarsalis*. These data indicate that *Wolbachia* infection is not present in *Cx. tarsalis* mosquitoes and the observed rare positive samples were due to sample misidentification or potentially contamination. The two *Cx. quinquefasciatus* samples were removed from further downstream analysis.

For fungal communities, *Ascomycota* and *Basidiomycota* were the two most abundant phyla representing over 79% of total abundance (Fig 5, Additional file 1). While *Basidomycota* was mainly composed of *Agaricomyctetes* (21.4%), *Ascomycota* was mostly represented by *Dothideomycetes* (53.1%) at the class level. Within *Dothidemycetes*, *Capnodiales* and *Pleosporales* were the two most abundant taxa at the order level representing over 53% of total abundance. *Mycosphaerella* (28.4%) and *Cladosporium* (16%) were the most abundant within *Dothideomycetes* while *Alternaria* was most abundant (3.8%) within *Pleosporales* at the genus level, and these three taxa were detected in 83 to 94% of all 72 samples. While *Ganoderma* was the second most abundant taxa (21.4%) belonging to *Basidiomycota*, its prevalence was relatively lower (36%).

**Fig. 5.**
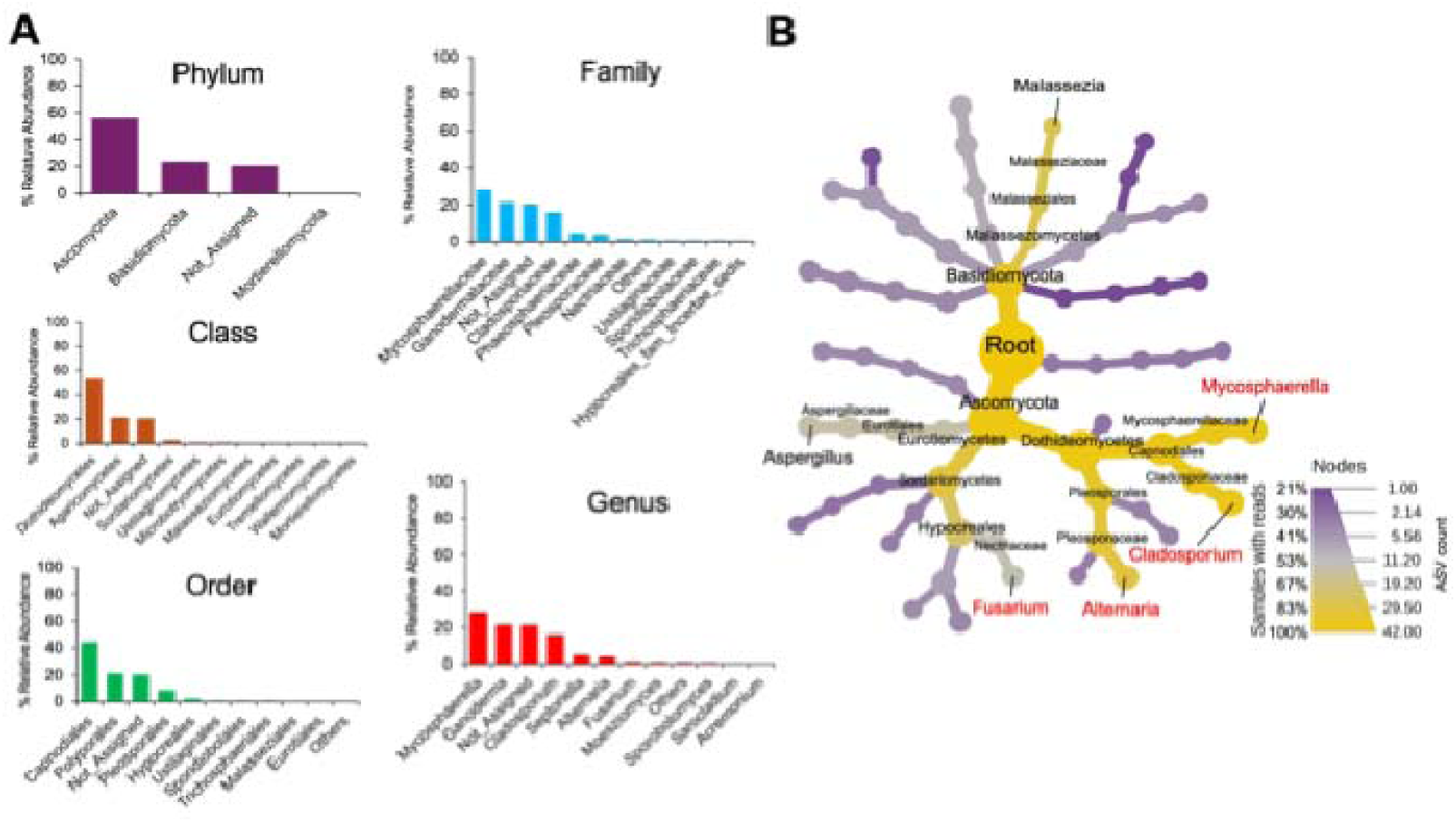
Fungal communities in *Culex tarsalis* mosquitoes. **A:** Percent relative abundance of the top 10 taxa. **B:** Heat tree showing hierarchical structure of representative taxa. Taxa with 50% prevalence or higher are labeled. Genera within the top 10 relative abundance (see Fig 2 A) are in red text. Color and node size indicate prevalence (% sequence read detected in all samples) and number of ASV, respectively. See Additional file 1 for heat tree with all taxa presented.

### Clustering of mosquito populations by microbial community structure

We identified a pattern where specific bacterial taxa at the genus level were more abundant in certain clusters of populations. Based on visual inspection and geographic proximity, we hypothesized three clustering groups of mosquito populations; CA1, CA2, CO1, CO2, CO3, TX1, and TX2 for group CCT (initials of California, Colorado and Texas), CA3, CA4, CA5, and CA6 for group CA (California), and UT1, UT2, UT3, WA1, WA2, and WA3 in group UW (initials of Utah and Washington) (Fig. 6A and Additional file 2). With this grouping, *Ralstonia* and *Pseudomonas* were the most abundant taxa in group CCT (70.2% relative abundance) and UW (66%), respectively, while *Zymobacter* (35.6%) and *Providencia* (31.3%) were most abundant in group CA (Additional file 2, 3).

**Fig. 6.**
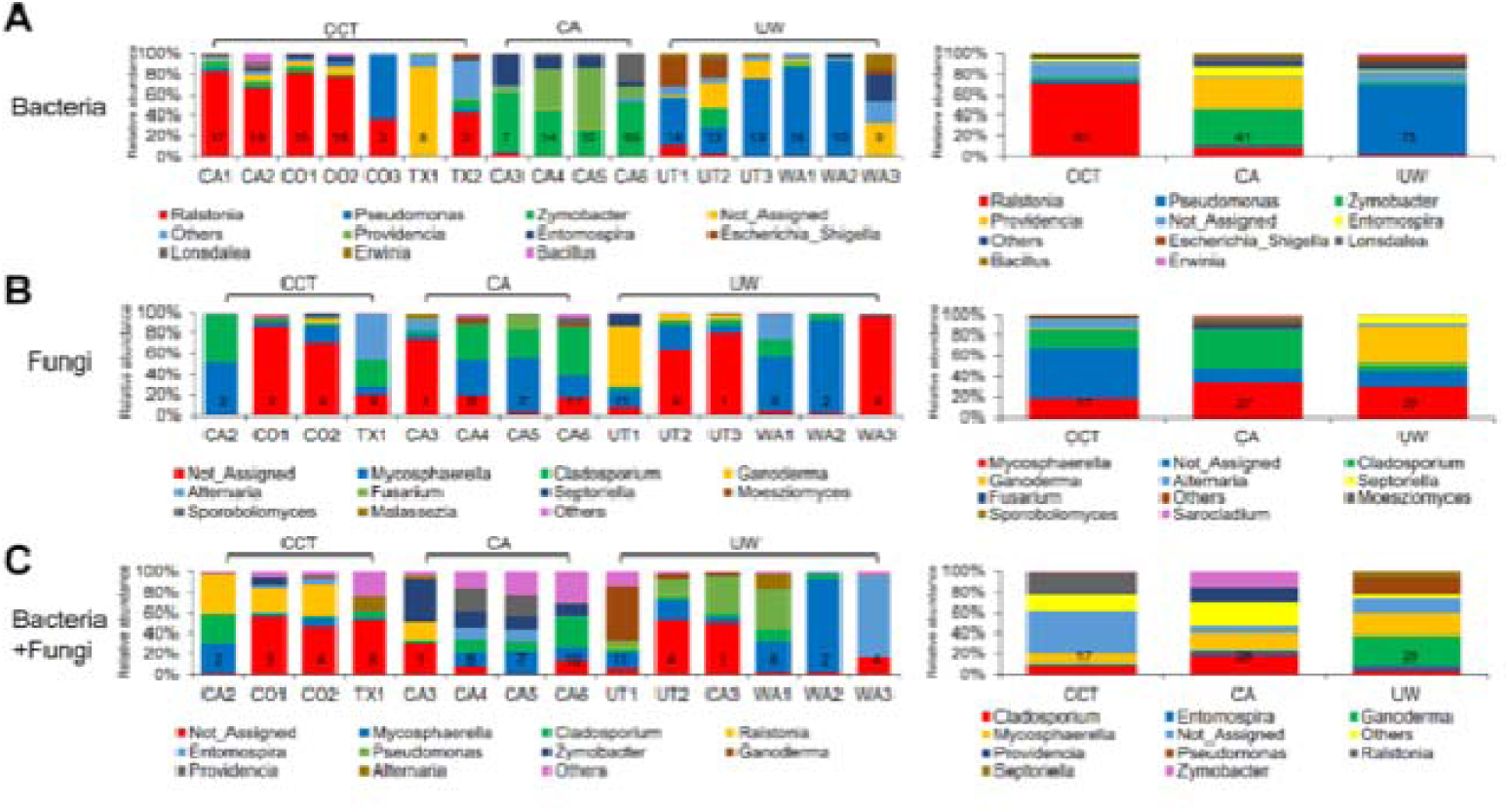
Hypothesized clustering pattern of mosquito populations based on community structure of **A:** bacteria, **B:** fungi, and **C:** bacteria/fungi combined at the population (left) or group level (right). Numbers within each bar graph indicate the number of mosquito samples assayed.

Clustering of fungal community structure according to this grouping was less pronounced (Fig. 6B; Additional file 2, 4). For example, while *Ganoderma* was most abundant in UW (36.5% relative abundance) and *Cladosporium* was relatively more abundant in CA (37.8%) than CCT (17.8%) or UW (6.3%), *Mycosphaerella* were found in all three groups with similar abundance ranging from 18.3 to 33.1%. Similarly to the case of bacteria, grouping was more pronounced at order or lower taxonomic levels (Additional file 4). When community structure of bacteria and fungi were combined, clustering was moderate, similarly to the case of fungal community (Fig 6C; Additional file 2; Additional file 5) as relative abundance of major taxa identified above (i.e., *Pseudomonas*, *Zymobacter*, *Providencia*, *Mycosphaerella*, *Cladosporium* and *Ganoderma*) followed the hypothesized grouping model.

### Alpha and Beta diversity, and Random Forest classification

Alpha diversity analyses, Chao1 (species richness) and Shannon (species diversity) indices were compared for community structure of bacteria, fungi, or bacterial/fungi combined (Fig. 7A). Chao1 indices were significantly different among the three groups (*P* < 0.05, Kruskal-Wallis) with higher index value in group CCT relative to CA (*P* = 0.006, Bonferroni corrected after pair-wise comparison among three groups) while Shannon indices were not different among three groups (*P* > 0.05, Kruskal-Wallis). For fungal communities, Chao1 indices were similar among three groups while a difference was observed for Shannon indices among the groups (*P* < 0.01, Kruskal-Wallis). When bacteria/fungi were combined, no difference was observed for Chao1 indices among the groups (*P* > 0.05, Kruskal-Wallis), but Shannon index was lowest for UW (*P* < 0.001, Kruskal-Wallis; Bonferroni corrected after pairwise comparison). Rarefaction curve analyses and comparison of sequencing depth among three groups indicated overall high sequencing depth across samples with minimal heteroskedasticity regardless of data filtering (Additional file 6). Beta diversity was next explored using Nonmetric Multidimensional Scaling (NMDS), revealing a significant differences in community structure among the three groups (PERMANOVA, *P* < 0.01) for bacterial, fungal, and bacterial/fungal combined (Fig. 7B).

**Fig. 7.**
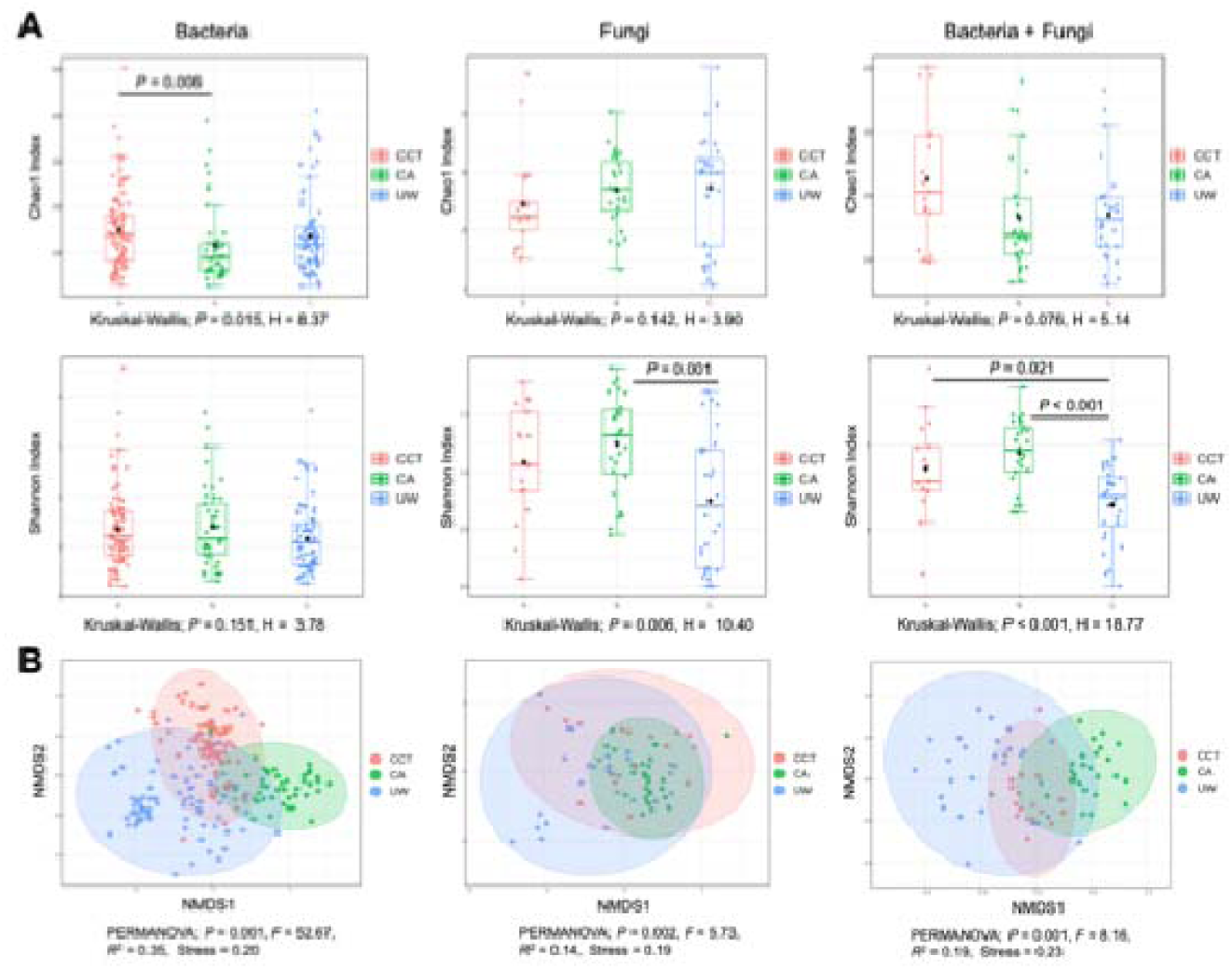
Alpha and Beta diversity analysis. **A:** Chao1 (species richness) and Shannon index (species diversity) were compared for Alpha diversity analysis. Pair-wise Mann-Witney tests were conducted between groups followed by Benjamini-Hochberg correction. **B:** Nonmetric Multidimensional Scaling (NMDS) with Bray-Curtis distance matric was estimated followed by Permutational Multivariate Analysis of Variance (PERMANOVA) for Beta diversity analysis.

We then performed machine learning based Random Forest classification analyses to validate our grouping model (Fig. 8). For bacterial communities, the mean Out-of-bag (OOB) error was less than 0.1 in five consecutive runs (Fig. 8A; Additional file 7) indicating that about 90% of predictions were correct by our grouping model, and *Ralstonia* was identified to have the greatest influence on the classification model followed by *Providencia*, *Pseudomonas*, *Collinsella* and *Zymobacter* (Fig. 8B). Similarly, dendrogram analysis largely supported our clustering model regardless of algorithms applied (Additional file 8). Classification was less clear for fungal data with slightly higher OOB error (0.228) (Fig. 8A; Additional file 7, 9). The top five fungal genera that had greatest influence on the classification were *Sporobolomyces*, *Sarocladium*, *Ganoderma*, *Cladosporium*, and *Aspergillus*; *Mycosphaerella* (the most abundant fungus) had relatively little influence (Fig. 8B). For combined bacterial/fungal data, intermediate OOB error (0.152) was observed (Fig. 8A; Additional file 7, 10), and *Zymobacter* had greatest influence on classification followed by *Providencia*, *Sporobolomyces*, *Pseudomonas*, and *Saccharibacter* (Fig. 8B).

**Fig. 8.**
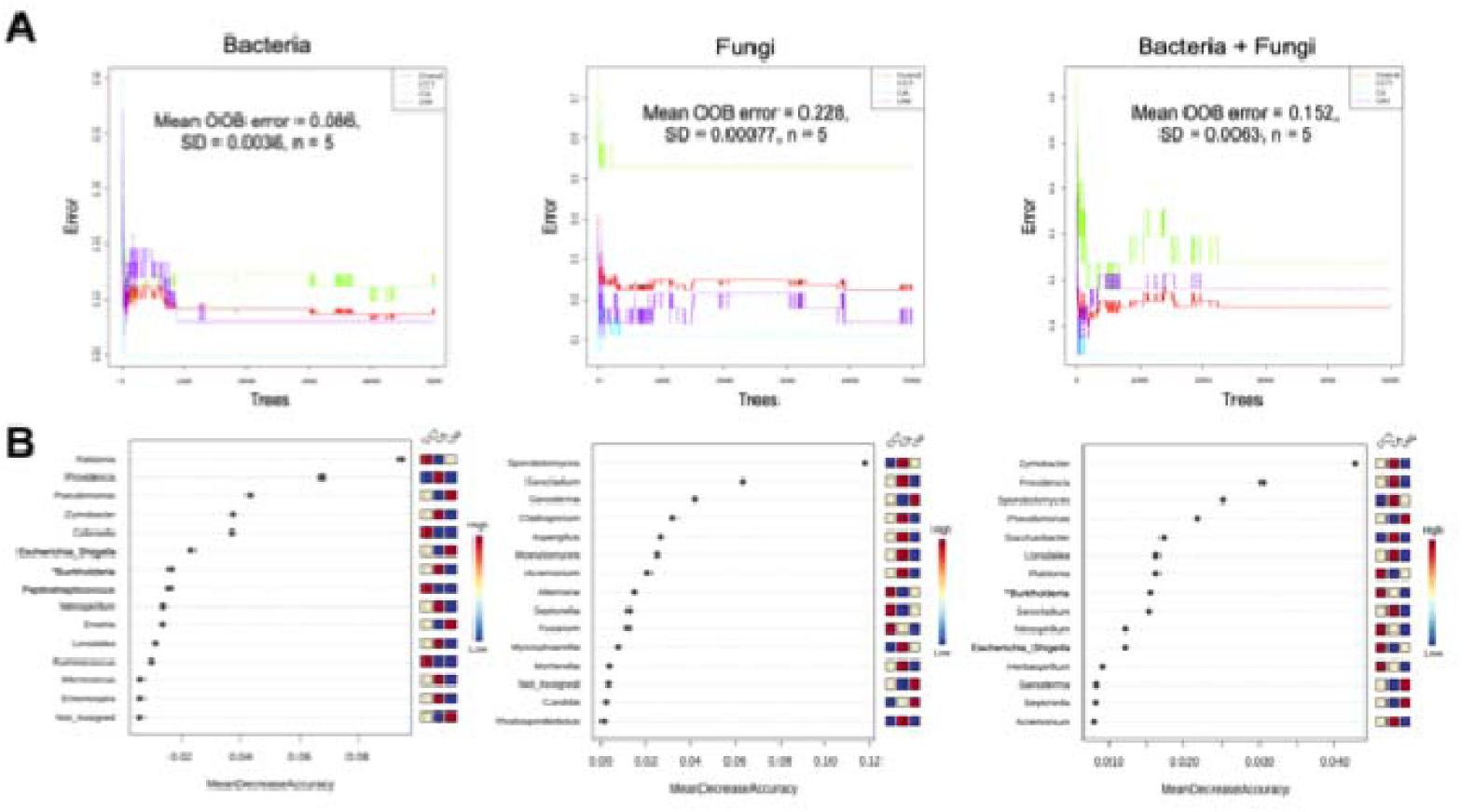
Random Forest classification analysis with **A:** representative tree plots, and **B:** top 15 taxa at genus level that had greatest influence on classification models. Mean OOB (Out Of Bag) errors were calculated using five consecutively generated OOB tables (see Additional file 7) (*,*Burkholderia, Caballeronia, Paraburkholderia*).

### Core microbiome, taxon differentiation, and correlation network

Prevalence (% detected in samples) profiles were characterized using Core microbiome analysis (Fig. 9). For bacterial data, *Ralstonia* was most prevalent followed by *Pseudomonas*, *Zymobacter*, *Entomospria* and *Escherichia* in all samples. When each group was analyzed separately, *Ralstonia* and *Pseudomonas* were most prevalent in group CCT and UW, respectively while *Zymobacter* and *Providencia* were most prevalent in group CA. For fungal data, *Cladosporium* and *Mycosphaerella* were widely found in all three groups while *Alternaria* and *Ganoderma* were found in CCT and UW respectively. A similar pattern was observed when bacteria/fungal data were combined.

**Fig. 9.**
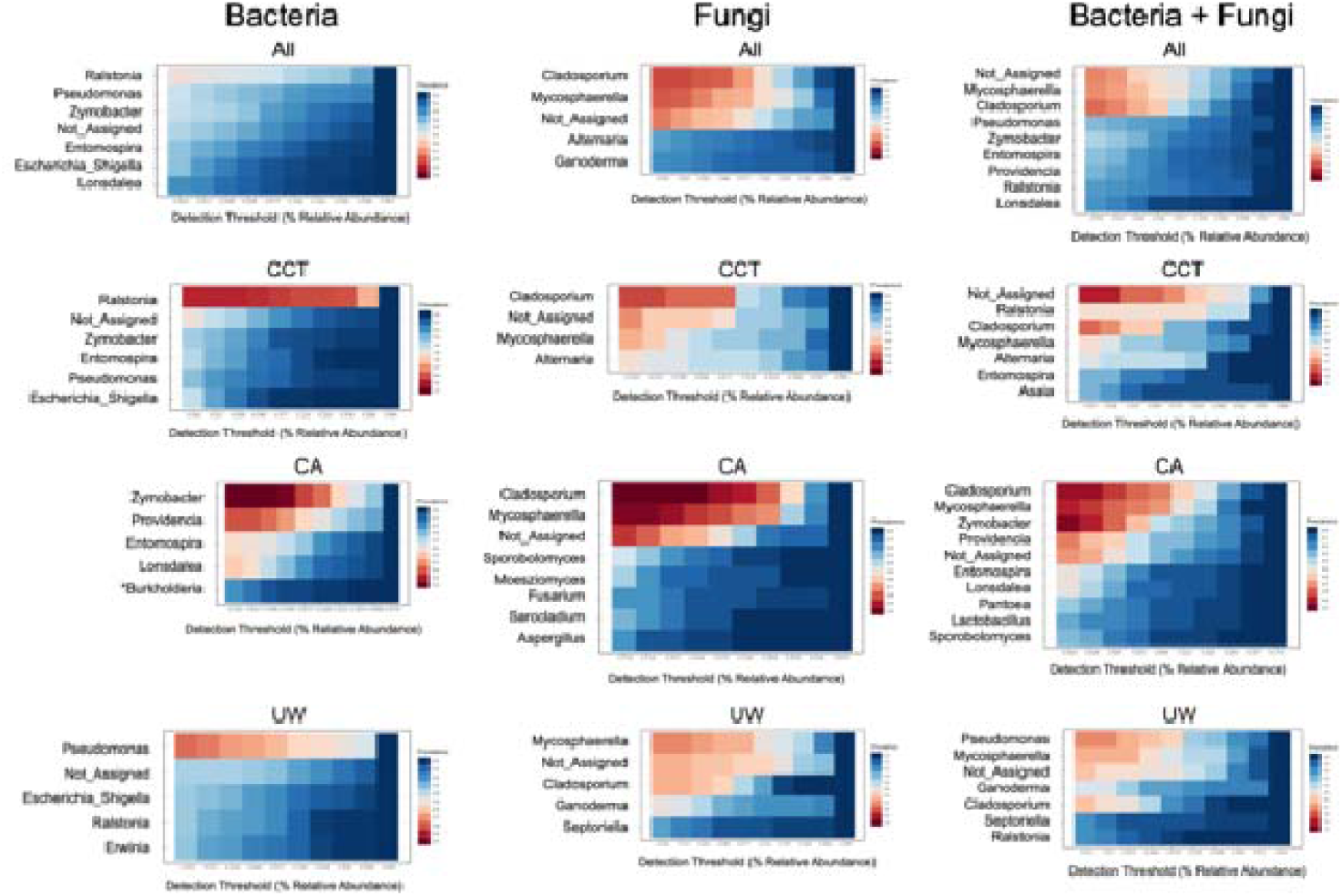
Core microbiome analysis. Taxa with 20% or higher prevalence are presented at genus level. The detection threshold was set at 0.01% relative abundance (*, *Burkholderia, Caballeronia, Paraburkholderia*).

The major taxa identified in Core microbiome analyses were consistently observed in Linear Discriminant Analysis with effect size (LEfSe) for bacterial data (Additional File 11). *Ralstonia* and *Pseudomonas* were significantly more abundant in CCT and UW, respectively while *Zymobacter* and *Providencia* were more abundant in group CA. However, for fungal data, *Cladosporium* and *Ganoderma* were more abundant in group CA and UW, respectively, but *Mycospharella* was not significantly enriched in either of three groups. A similar pattern was observed for combined bacterial/fungal data.

To explore whether interactions among taxa had influence on our clustering model, we investigated correlation network for the major taxa identified above. For bacteria, *Ralstonia*, *Pseudomonas*, and *Zymobacter* were uncorrelated (Additional file 12; Additional file 13) while for fungi, *Mycosphaerella* and *Cladosporium* were positively correlated, while *Mycosphaerella* was negatively correlated with *Alternaria*. In total, 33 significant correlations were identified between bacteria and fungi, and 85% (28 cases) of them were negative. The two most abundant fungi, *Mycosphaerella* and *Cladosporium,* accounted for 50% (14 cases) of total negative correlations (Additional file 14A). Consistent with this result, species richness (Observed ASV/Chao1) was negatively correlated between bacteria and fungi (*R* = −0.3, *P* = 0.0119) (Additional file 14B). These results indicate a possible exclusive interaction between bacteria and fungi within the mosquito microbiota.

### Relationship between environmental factors and community structure

Because *Cx. tarsalis* population genetic structure was recently observed to be correlated with environmental factors [6], we explored whether environmental factors were correlated with microbial community structure or species richness/diversity in this mosquito. We first examined the relationship between environmental factors (e.g., longitude, latitude, temperature, and precipitation) and Alpha diversity indices (Fig. 10A). For bacteria, ASV/Chao1 and ACE were positively correlated with longitude and precipitation while these two indices were negatively correlated with temperature. For fungi, ASV/Chao1 and ACE were negatively correlated with longitude while these two indices together with Shannon and Simpson indices were positively correlated with temperature. Latitude was also positively correlated with Shannon and Simpson indices. For combined bacterial/fungal data, no correlation was found with longitude for any indices, but all five indices were positively correlated with temperature and negatively correlated with latitude. These results suggest species richness/diversity is shaped by multiple environmental factors. Particularly, species richness of bacteria and fungi were negatively correlated along longitude.

**Fig. 10.**
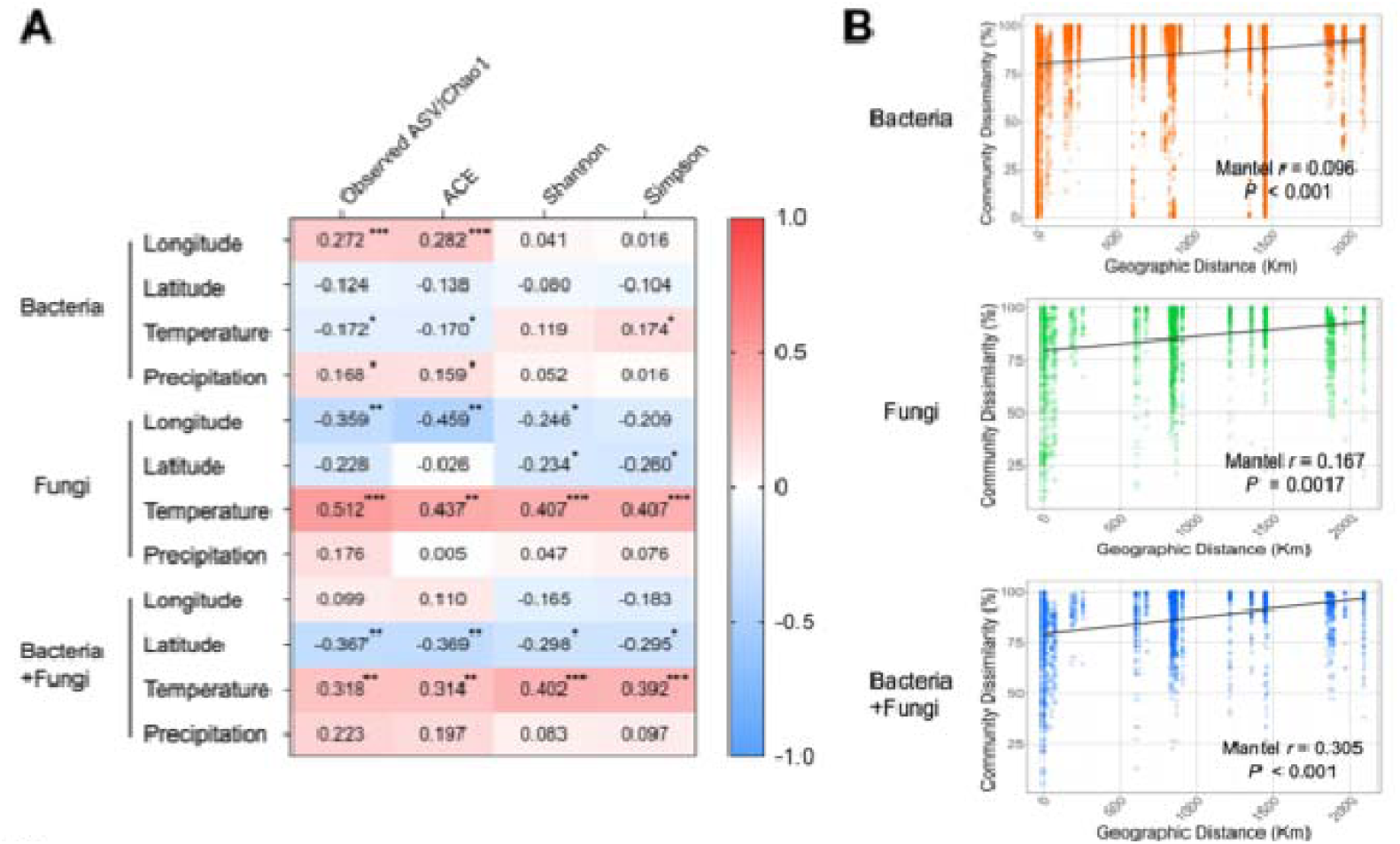
Correlation analyses between environmental factors and species richness/diversity or geographic structure. **A:** Correlation between abiotic environmental factors and Alpha diversity indices using linear regression. Positive and negative numbers indicate positive (red) and negative (blue) correlation, respectively (***, *P* < 0.001; **, *P* < 0.01; *, *P* < 0.05). **B:** Correlation between geographic distance and Bray-Curtis dissimilarity index using Mantel test. Line plots with shaded areas indicate linear regression with 95% CI.

We noted a significant positive correlation between geographic distance and community dissimilarity for bacterial, fungal and combined bacteria/fungal data (Mantel test, *P* < 0.01; Fig. 10B), indicating that the community structure is more dissimilar as distance between collection sites increases for both bacteria and fungi and suggesting possible phylogenomic signal of microbial diversity in *Cx. tarsalis*.

## Discussion

Our population genetic analysis of *Cx. tarsalis* across the western USA identified essentially the same population structure that previous microsatellite and individual RADSeq analyses. In total, these analyses demonstrate that gene flow is substantial between *Cx. tarsalis* populations across wide geographic distances, and that the metapopulation structure can be defined in terms of 3-4 large metapopulation clusters encompassing the Midwest, the Pacific Coast, the Sonoran desert, and potentially the Northwest [1,3,6]. Although it does not resolve individuals, PoolRADSeq produced similar population-level results at significantly lower cost.

There are no previous studies on the diversity of fungi colonizing *Cx. tarsalis*, and previous bacterial microbiome studies in this species were limited to analysis of mosquitoes that were reared in artificial field mesocosms [29]. These mesocosms were constructed in an attempt to mimic the natural habitat of this mosquito; however, the resulting microbial composition data differed significantly compared to our results using true wild-caught *Cx. tarsalis*. The bacterial microbiomes of mesocosm-reared mosquitoes were dominated by *Thorsellia* [29] a common aquatic bacterial taxon that has been previously associated with mosquitoes and other aquatic species [30, 41], and likely reflects the unnatural microbial colonization of the mesocosm habitats. In our study, we observed that distinct microbial taxa were enriched in wild-reared mosquitoes, and these taxa differed depending on the geographic region the mosquitoes were collected from. These data suggest that microbiome studies should, if at all possible, be conducted on individuals that were collected from natural habitats, as artificial habitats may not reflect the microbial diversity observed in nature and can give misleading results.

*Wolbachia* is a maternally inherited alphaproteobacterial symbiont that is of interest as an agent for control of arbovirus transmission in mosquitoes [27,28,29,40,43,44]. Previous studies suggested that *Cx*. *tarsalis* is not naturally infected with *Wolbachia* [39]. Artificial transfection with *Wolbachia* in *Cx*. *tarsalis* was shown to increase, rather than decrease, their susceptibility to WNV, thus detection of *Wolbachia* in this species has significant implication for the epidemiology of WNV transmission (and potentially other arboviruses) [45]. Therefore, when we detected *Wolbachia* by 16S amplicon sequencing in a small subset of individuals we attempted to independently validate the finding. Out of 12 samples where *Wolbachia* was detected by amplicon sequencing, we were able to independently validate the infection in only two individuals, which were ultimately molecularly identified as *Cx*. *quinquefasciatus;* a species which is known to be natively infected with *Wolbachia* [39,40]. In the remaining 10 samples, *Wolbachia* infection could not be verified, and these samples were molecularly identified as *Cx*. *tarsalis*. In total, these data affirm previous findings that *Cx*. *tarsalis* is not naturally infected with *Wolbachia* and highlight the importance of independently verifying low frequency *Wolbachia* infections detected solely on the basis of 16S amplicon microbiome sequencing.

Recent studies identified correlations between *Cx. tarsalis* population genetics and environmental variables such as temperature, humidity, and vegetation [6]. We therefore examined potential links between microbiome composition and environment. We found that bacterial diversity was positively correlated with longitude and precipitation, and negatively correlated with temperature. Fungal diversity was negatively correlated with longitude and positively correlated with temperature. The mechanisms underlying these relationships are at present unclear, but the data suggest that correlations between environmental parameters and mosquito genetics are confounded by microbial interactions. Future studies are indicated to tease apart the mechanisms underlying these interactions.

Our data suggests that microbiome structure in *Cx. tarsalis* is, to some extent, correlated with population genetic structure (Fig 11). While this relationship is not completely one-to-one with the genetic clusters defined by RADSeq, it is significant and partially overlaps with these clusters. Mantel regression demonstrates significant isolation by distance in microbial structure. Phylogenomic signal is more pronounced for bacterial data compared to fungi. Generally, mosquitoes acquire bacterial microbiome associates from the aquatic larval environment [19,46]. Less is known about how mosquitoes acquire fungal associates. Differences between bacteria and fungi could reflect different modes of acquisition between these two groups, differences in retention between bacteria vs. fungi during mosquito development, or differences in mosquito immune response to bacteria vs. fungi in response to colonization.

**Fig. 11.**
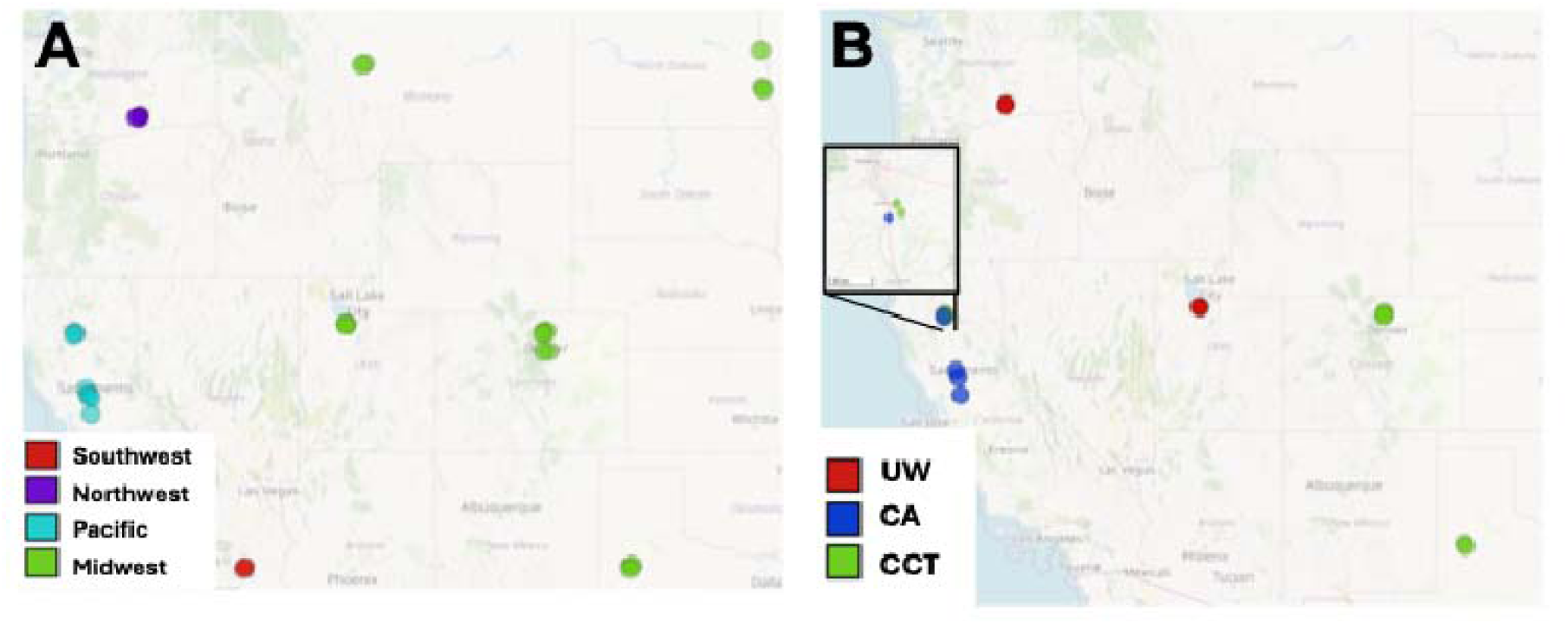
Comparison of **A:** *Cx*. *tarsalis* population genetic structure and **B:** Microbiome community structure. Colors denote grouping identities as described in the text.

## Conclusions

Microbial associates have been demonstrated to affect pathogen infection and transmission in a variety of mosquito-pathogen systems [15,16,17,18,19,20,21,22, 23,24,25,26,27,28]. This is the first study to examine diversity of microbial association in natural populations of *Cx. tarsalis* across its species range. Identification of microbial taxa that can affect WNV or other pathogens in this mosquito has significant implications for understanding the spread and invasion of the viral pathogens transmitted by this mosquito across space and time.

## Acknowledgements

We thank our collaborators for providing mosquito samples used in this analysis, including Anna Wanek, Broox Boze, Jason Williams, Nikki Harris and Will Schlatmann (Vector Disease Control International), Conlin Reis (Fresno Westside MAD), Corey Brelsfoard (Texas Tech University), Gary Hatch (Mosquito Abatement District-Davis), Jared Larmirante (Cass County Vector Control), Jeff Hamik (Nebraska Department of Health and Human Services), John Albright (Shasta Mosquito and Vector Control District), Joshua Blystone (Cascade County Weed & Mosquito Division), Kevin Shoemaker (Benton County Mosquito Control District), Kim Hung (Coachella Valley Mosquito & Vector Control District), Ryan Smith (Iowa State University), Sarah Wheeler (Sacramento-Yolo Mosquito & Vector Control District), Scott Bradshaw (Tooele Valley Mosquito Abatement District), Scott Larson (Metropolitan Mosquito Control District), and Todd Hanson (Mosquito Control Division, Grand Forks Public Health Department).

## Author contributions

All authors contributed intellectually to and agreed to this submission. ES and JLR designed the experiments. ES (and others acknowledged above) collected field mosquito samples. ES, TW, and NN conducted experiments and collected the data. ES, NH, and JPS analyzed the data. ES and JLR wrote the manuscript, and NH, TW, JPS, and NN provided feedback. The authors read and approved the final manuscript.

## Availability of data and materials

Raw bacterial microbiome data, fungal microbiome data, and PoolRADseq data are available at zenodo.org/records/14081783, zenodo.org/records/14080098, and zenodo.org/records/14081214.

## Code availability

Code for analyses is available at zenodo.org/records/17014950.

## Ethics approval and consent to participate

Not applicable.

## Consent for publication

Not applicable.

## References

1. Venkatesan M, Rasgon JL. Population genetic data suggest a role for mosquito-mediated dispersal of West Nile virus across the western United States. Mol Ecol. 2010; 19:1573–84.

2. Reisen WK. The contrasting bionomics of Culex mosquitoes in western North America. J Am Mosq Control Assoc. 2012; 28(4 Suppl):82-91.

3. Venkatesan M, Westbrook CJ, Hauer MC, Rasgon JL. Evidence for a population expansion in the West Nile virus vector Culex tarsalis. Mol Biol Evol. 2007; 24:1208–18.

4. Reisen WK, Lothrop HD, Hardy JL. Bionomics of Culex tarsalis (Diptera: Culicidae) in relation to arbovirus transmission in southeastern California. J Med Entomol. 1995; 32:316–27.

5. Goddard LB, Roth AE, Reisen WK, Scott TW. Vector competence of California mosquitoes for West Nile virus. Emerg Infect Dis. 2002; 8:1385–91.

6. Liao Y, Islam T, Noorai R, Streich J, Saski C, Cohnstaedt LW, Cooper EA. Climate Adaptation and Genetic Differentiation in the Mosquito Species Culex tarsalis. Genome Biol Evol. 2025; 17:evaf143.

7. Hugo LE, Monkman J, Dave KA, Wockner LF, Birrell GW, Norris EL, Kienzle VJ, Sikulu MT, Ryan PA, Gorman JJ, Kay BH. Proteomic biomarkers for ageing the mosquito Aedes aegypti to determine risk of pathogen transmission. PLoS One. 2013;8:e58656. doi: 10.1371/journal.pone.0058656.

8. Hardy JL, Apperson G, Asman SM, Reeves WC. Selection of a strain of Culex tarsalis highly resistant to infection following ingestion of western equine encephalomyelitis virus. Am J Trop Med Hyg. 1978; 27(2 Pt 1):313-21.

9. Bosio CF, Beaty BJ, Black WC 4th. Quantitative genetics of vector competence for dengue-2 virus in Aedes aegypti. Am J Trop Med Hyg. 1998; 59:965-70.

10. Black WC 4th, Bennett KE, Gorrochótegui-Escalante N, Barillas-Mury CV, Fernández-Salas I, de Lourdes Muñoz M, Farfán-Alé JA, Olson KE, Beaty BJ. Flavivirus susceptibility in Aedes aegypti. Arch Med Res. 2002; 33:379-88.

11. Bennett KE, Flick D, Fleming KH, Jochim R, Beaty BJ, Black WC 4th. Quantitative trait loci that control dengue-2 virus dissemination in the mosquito Aedes aegypti. Genetics. 2005; 170:185-94.

12. Alto BW, Bettinardi D. Temperature and dengue virus infection in mosquitoes: independent effects on the immature and adult stages. Am J Trop Med Hyg. 2013; 88:497–505.

13. Bosio CF, Fulton RE, Salasek ML, Beaty BJ, Black WC 4th. Quantitative trait loci that control vector competence for dengue-2 virus in the mosquito Aedes aegypti. Genetics. 2000; 156:687-98.

14. Bennett KE, Olson KE, Muñoz Mde L, Fernandez-Salas I, Farfan-Ale JA, Higgs S, Black WC 4th, Beaty BJ. Variation in vector competence for dengue 2 virus among 24 collections of Aedes aegypti from Mexico and the United States. Am J Trop Med Hyg. 2002; 67:85-92.

15. Huang W, Rodrigues J, Bilgo E, Tormo JR, Challenger JD, De Cozar-Gallardo C, Pérez-Victoria I, Reyes F, Castañeda-Casado P, Gnambani EJ, Hien DFS, Konkobo M, Urones B, Coppens I, Mendoza-Losana A, Ballell L, Diabate A, Churcher TS, Jacobs-Lorena M. Delftia tsuruhatensis TC1 symbiont suppresses malaria transmission by anopheline mosquitoes. Science. 2023; 381:533-540.

16. Angleró-Rodríguez YI, Blumberg BJ, Dong Y, Sandiford SL, Pike A, Clayton AM, Dimopoulos G. A natural Anopheles-associated Penicillium chrysogenum enhances mosquito susceptibility to Plasmodium infection. Sci Rep. 2016; 6:34084.

17. Bahia AC, Dong Y, Blumberg BJ, Mlambo G, Tripathi A, BenMarzouk-Hidalgo OJ, Chandra R, Dimopoulos G. Exploring Anopheles gut bacteria for Plasmodium blocking activity. Environ Microbiol. 2014; 16:2980–94.

18. Hegde S, Rasgon JL, Hughes GL. The microbiome modulates arbovirus transmission in mosquitoes. Curr Opin Virol. 2015; 15:97–102.

19. Foo A, Brettell LE, Nichols HL; 2022 UW-Madison Capstone in Microbiology Students; Medina Muñoz M, Lysne JA, Dhokiya V, Hoque AF, Brackney DE, Caragata EP, Hutchinson ML, Jacobs-Lorena M, Lampe DJ, Martin E, Valiente Moro C, Povelones M, Short SM, Steven B, Xu J, Paustian TD, Rondon MR, Hughes GL, Coon KL, Heinz E. MosAIC: An annotated collection of mosquito-associated bacteria with high-quality genome assemblies. PLoS Biol. 2024; 22:e3002897.

20. Ramirez JL, Short SM, Bahia AC, Saraiva RG, Dong Y, Kang S, Tripathi A, Mlambo G, Dimopoulos G. Chromobacterium Csp_P reduces malaria and dengue infection in vector mosquitoes and has entomopathogenic and in vitro anti-pathogen activities. PLoS Pathog. 2014; 10:e1004398.

21. Pumpuni CB, Beier MS, Nataro JP, Guers LD, Davis JR. Plasmodium falciparum: inhibition of sporogonic development in Anopheles stephensi by gram-negative bacteria. Exp Parasitol. 1993; 77:195–9.

22. Mourya DT, Gokhale MD, Pidiyar V, Barde PV, Patole M, Mishra AC, Shouche Y. Study of the effect of the midgut bacterial flora of Culex quinquefasciatus on the susceptibility of mosquitoes to Japanese encephalitis virus. Acta Virol. 2002; 46:257–60.

23. Cirimotich CM, Dong Y, Clayton AM, Sandiford SL, Souza-Neto JA, Mulenga M, Dimopoulos G. Natural microbe-mediated refractoriness to Plasmodium infection in Anopheles gambiae. Science. 2011;332:855–8.

24. Apte-Deshpande A, Paingankar M, Gokhale MD, Deobagkar DN. Serratia odorifera a midgut inhabitant of Aedes aegypti mosquito enhances its susceptibility to dengue-2 virus. PLoS One. 2012; 7:e40401.

25. Sun X, Wang Y, Yuan F, Zhang Y, Kang X, Sun J, Wang P, Lu T, Sae Wang F, Gu J, Wang J, Xia Q, Zheng A, Zou Z. Gut symbiont-derived sphingosine modulates vector competence in Aedes mosquitoes. Nat Commun. 2024; 15:8221.

26. Garza-Hernández JA, Rodríguez-Pérez MA, Salazar MI, Russell TL, Adeleke MA, de Luna-Santillana Ede J, Reyes-Villanueva F. Vectorial capacity of Aedes aegypti for dengue virus type 2 is reduced with co-infection of Metarhizium anisopliae. PLoS Negl Trop Dis. 2013; 7:e2013.

27. Dutra HL, Rocha MN, Dias FB, Mansur SB, Caragata EP, Moreira LA. Wolbachia Blocks Currently Circulating Zika Virus Isolates in Brazilian Aedes aegypti Mosquitoes. Cell Host Microbe. 2016; 19:771–4.

28. Hugo LE, Rašić G, Maynard AJ, Ambrose L, Liddington C, Thomas CJE, Nath NS, Graham M, Winterford C, Wimalasiri-Yapa BMCR, Xi Z, Beebe NW, Devine GJ. Wolbachia wAlbB inhibit dengue and Zika infection in the mosquito Aedes aegypti with an Australian background. PLoS Negl Trop Dis. 2022; 16:e0010786.

29. Duguma D, Hall MW, Rugman-Jones P, Stouthamer R, Terenius O, Neufeld JD, Walton WE. Developmental succession of the microbiome of Culex mosquitoes. BMC Microbiol. 2015; 24;15:140.

30. Kämpfer P, Lindh JM, Terenius O, Haghdoost S, Falsen E, Busse HJ, Faye I. Thorsellia anophelis gen. nov., sp. nov., a new member of the Gammaproteobacteria. Int J Syst Evol Microbiol. 2006; 56(Pt 2):335–338.

31. Fu X, Dou J, Mao J, Su H, Jiao W, Zhang L, Hu X, Huang X, Wang S, Bao Z. RADtyping: an integrated package for accurate de novo codominant and dominant RAD genotyping in mapping populations. PLoS One. 2013; 8:e79960.

32. Callahan BJ, McMurdie PJ, Rosen MJ, Han AW, Johnson AJ, Holmes SP. DADA2: High-resolution sample inference from Illumina amplicon data. Nat Methods. 2016; 13:581–3.

33. Bokulich NA, Kaehler BD, Rideout JR, Dillon M, Bolyen E, Knight R, et al. Optimizing taxonomic classification of marker-gene amplicon sequences with QIIME 2 ’ s q2-feature-classifier plugin. Microbiome. 2018; 6:17.

34. Bolyen E, Rideout JR, Dillon MR, Bokulich N, Abnet CC, Al-Ghalith GA, et al. Reproducible, interactive, scalable and extensible microbiome data science using QIIME 2. Nat Biotechnol. 2019; 37:852–7.

35. Lu Y, Zhou G, Ewald J, Pang Z, Shiri T, Xia J. MicrobiomeAnalyst 2.0: comprehensive statistical, functional and integrative analysis of microbiome data. Nucleic Acids Res. 2023; 51:W310–W318.

36. Zhou W, Rousset F, O’Neil S. Phylogeny and PCR-based classification of Wolbachia strains using wsp gene sequences. Proc Biol Sci. 1998; 265:509–15.

37. Rasgon JL, Cornel AJ, Scott TW. Evolutionary history of a mosquito endosymbiont revealed through mitochondrial hitchhiking. Proc Biol Sci. 2006; 273:1603–11.

38. Rasgon JL, Scott TW. An initial survey for Wolbachia (Rickettsiales: Rickettsiaceae) infections in selected California mosquitoes (Diptera: Culicidae). J Med Entomol. 2004; 41:255–7.

39. Rasgon JL, Scott TW. Wolbachia and cytoplasmic incompatibility in the California Culex pipiens mosquito species complex: parameter estimates and infection dynamics in natural populations. Genetics. 2003; 165:2029–38.

40. Chavshin AR, Oshaghi MA, Vatandoost H, Pourmand MR, Raeisi A, Terenius O. Isolation and identification of culturable bacteria from wild Anopheles culicifacies, a first step in a paratransgenesis approach. Parasit Vectors. 2014; 4;7:419.

41. Taha MME, Abdelwahab SI, Mohamed HY, Jerah A, Alabsi AM, Abdullah SM, Oraibi B, Alfaifi HA, Babiker YOH, Aziz Ibrahim IA, Alshahrani S, Farasani AM, Alamer AS, Altherwi T. Comprehensive review of Wolbachia research (1936-2024): Global landscape, mapping progress and themes. Parasite Epidemiol Control. 2025; 30:e00438.

42. Anders KL, Indriani C, Ahmad RA, Tantowijoyo W, Arguni E, Andari B, Jewell NP, Dufault SM, Ryan PA, Tanamas SK, Rancès E, O’Neill SL, Simmons CP, Utarini A. Update to the AWED (Applying Wolbachia to Eliminate Dengue) trial study protocol: a cluster randomised controlled trial in Yogyakarta, Indonesia. Trials. 2020; 21:429.

43. Ribeiro Dos Santos G, Durovni B, Saraceni V, Souza Riback TI, Pinto SB, Anders KL, Moreira LA, Salje H. Estimating the effect of the wMel release programme on the incidence of dengue and chikungunya in Rio de Janeiro, Brazil: a spatiotemporal modelling study. Lancet Infect Dis. 2022; 22:1587-1595.

44. Dodson BL, Hughes GL, Paul O, Matacchiero AC, Kramer LD, Rasgon JL. Wolbachia enhances West Nile virus (WNV) infection in the mosquito Culex tarsalis. PLoS Negl Trop Dis. 2014; 8:e2965.

45. Coon KL, Vogel KJ, Brown MR, Strand MR. Mosquitoes rely on their gut microbiota for development. Mol Ecol. 2014; 23:2727–39.

