## Supplementary material for "Hologenomic structure of bacterial and fungal community composition in the West Nile virus vector *Culex tarsalis*"

**Additional file legends**

**Additional File 1**. Heat trees showing hierarchical structure of all taxa.

**Additional File 2.** Hypothesized clustering of mosquito populations showing community structure in individual samples for bacteria, fungi and bacteria and fungi combined at genus level (unfiltered original data).

**Additional File 3.** Hypothesized clustering of mosquito populations showing bacterial community structure in individual samples at family or higher taxonomic levels (unfiltered original data).

**Additional File 4.** Hypothesized clustering of mosquito populations showing fungal community structure in individual samples at family or higher taxonomic levels (unfiltered original data).

**Additional File 5.** Hypothesized clustering of mosquito populations showing community structure of bacteria and fungi combined in individual samples at family or higher taxonomic levels (unfiltered original data).

**Additional File 6.** Rarefaction curves (line plots) and comparison of sequencing depth among three groups (dot plots) for original and filtered data. Significance was determined using Mann-Whitney U test with Bonferroni correction for sequencing depth comparison (*, $P < 0.05$). Unfiltered original data were used for Alpha diversity analysis while all other comparative analyses used filtered data.

**Additional File 7.** Out of Bag (OOB) error tables. Five OOB tables were consecutively generated to estimate mean OOB errors for Random Forest classification analysis.

**Additional File 8.** Dendrograms for bacterial communities generated with Bray -Curtis index using four different algorithms.

**Additional File 9.** Dendrograms for fungal communities generated with Bray -Curtis index using four different algorithms.

**Additional File 10.** Dendrograms for bacterial/fungal communities generated with Bray -Curtis index using four different algorithms.

**Additional File 11.** Linear discriminant analysis Effect Size (LEfSe). Top 15 significantly enriched taxa at genus level are presented ($P < 0.05$) (*, *Burkholderia_Caballeronia_Paraburkholderia*).

**Additional File 12.** Correlation network analysis. Color of node indicates phylum. Node size represents the abundance of the taxon. Red and blue lines indicate positive and negative correlation, respectively (numbers indicate strength of correlation). Correlations are highlighted for major taxa such as *Ralstonia*, *Pseudomonas*, and *Zymobacter*. Complete correlation information for the bacterial community is available in Additional file 13.

**Additional File 13.** Correlation network table for bacterial community.

**Additional File 14. A:** Correlation network analysis between bacterial and fungal communities. Only significant correlations are presented. Positive and negative numbers indicate positive (red) and negative (blue) correlations, respectively. **B:** Correlation analysis for Alpha diversity indices between bacteria and fungi using linear regression (dotted lines indicate 95% CI for linear regression line). Each dot represents an individual mosquito sample.

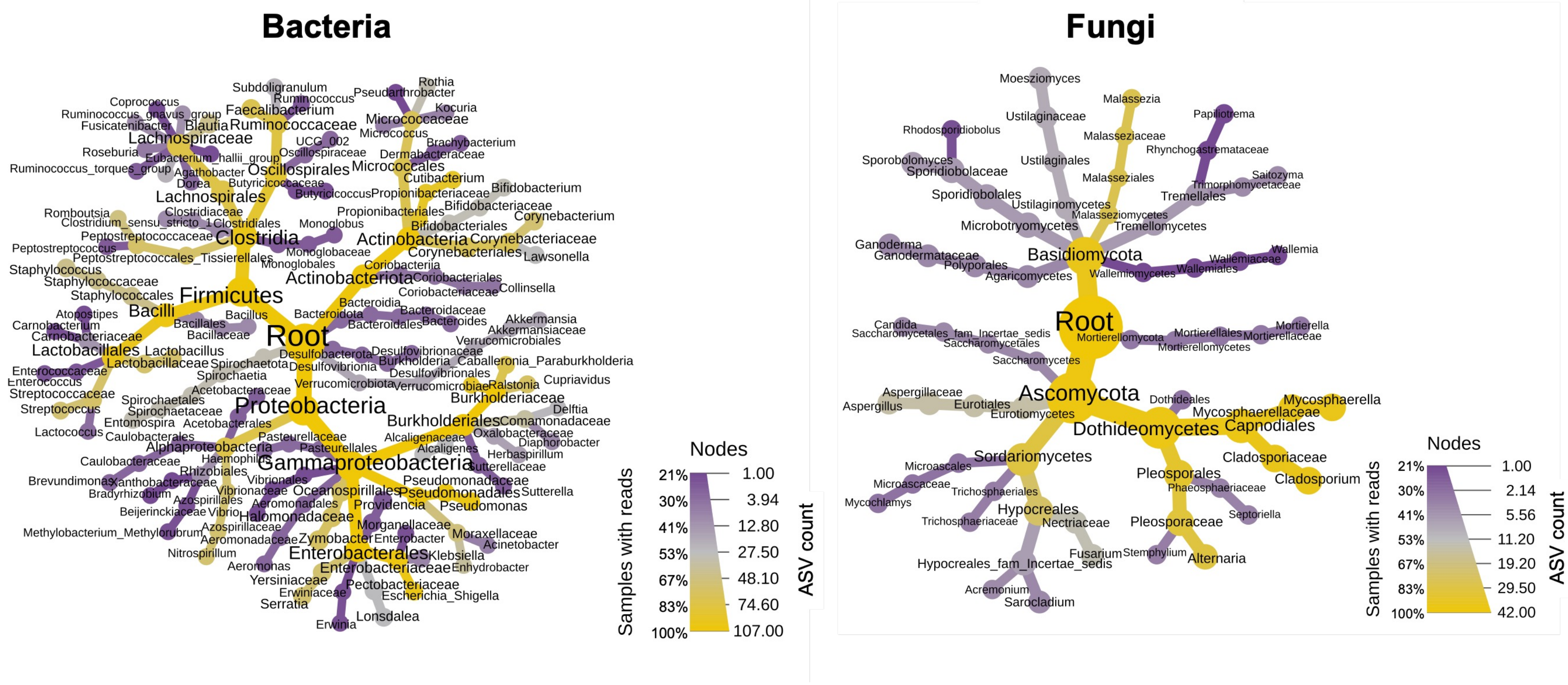


Additional file 1

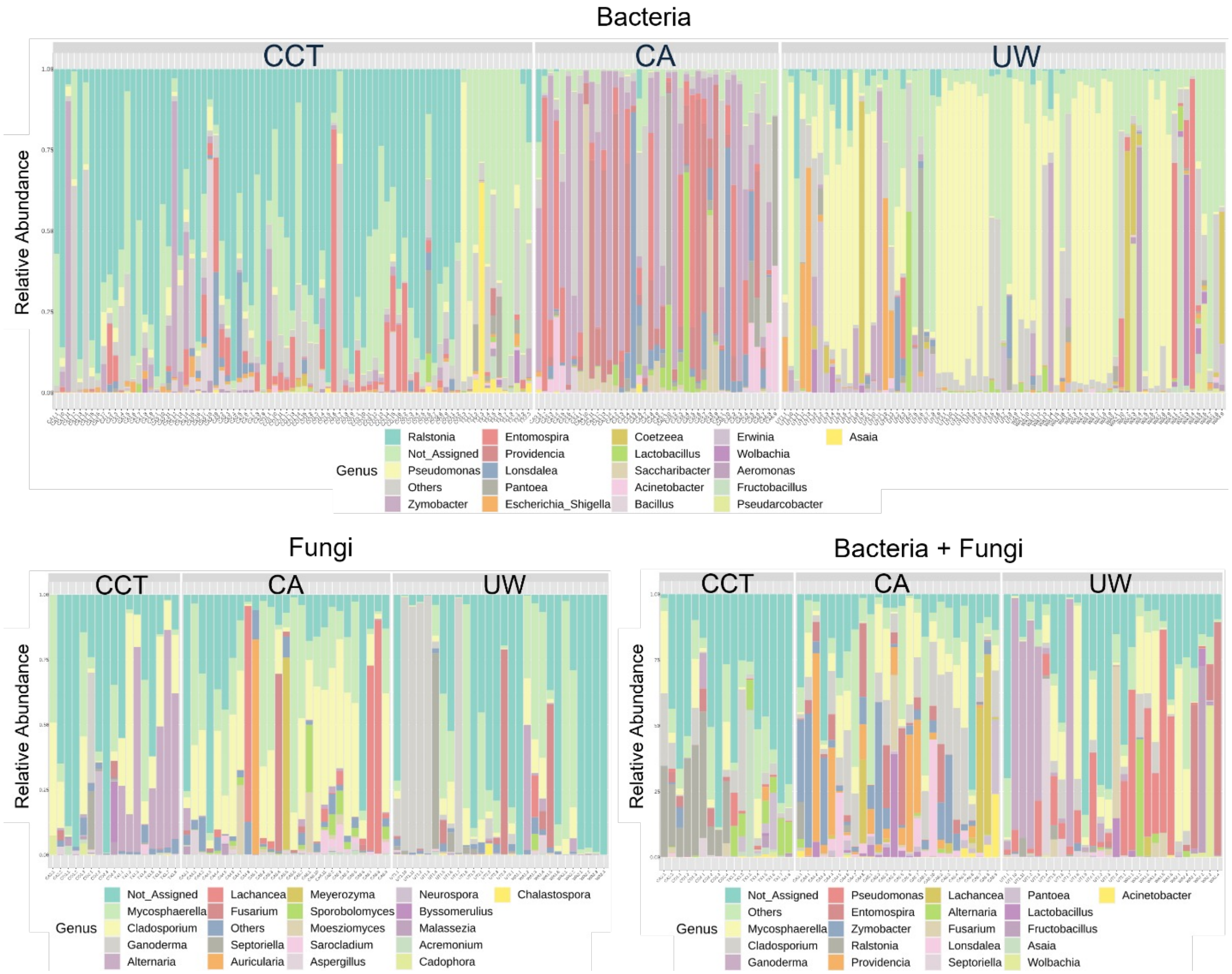


Additional file 2

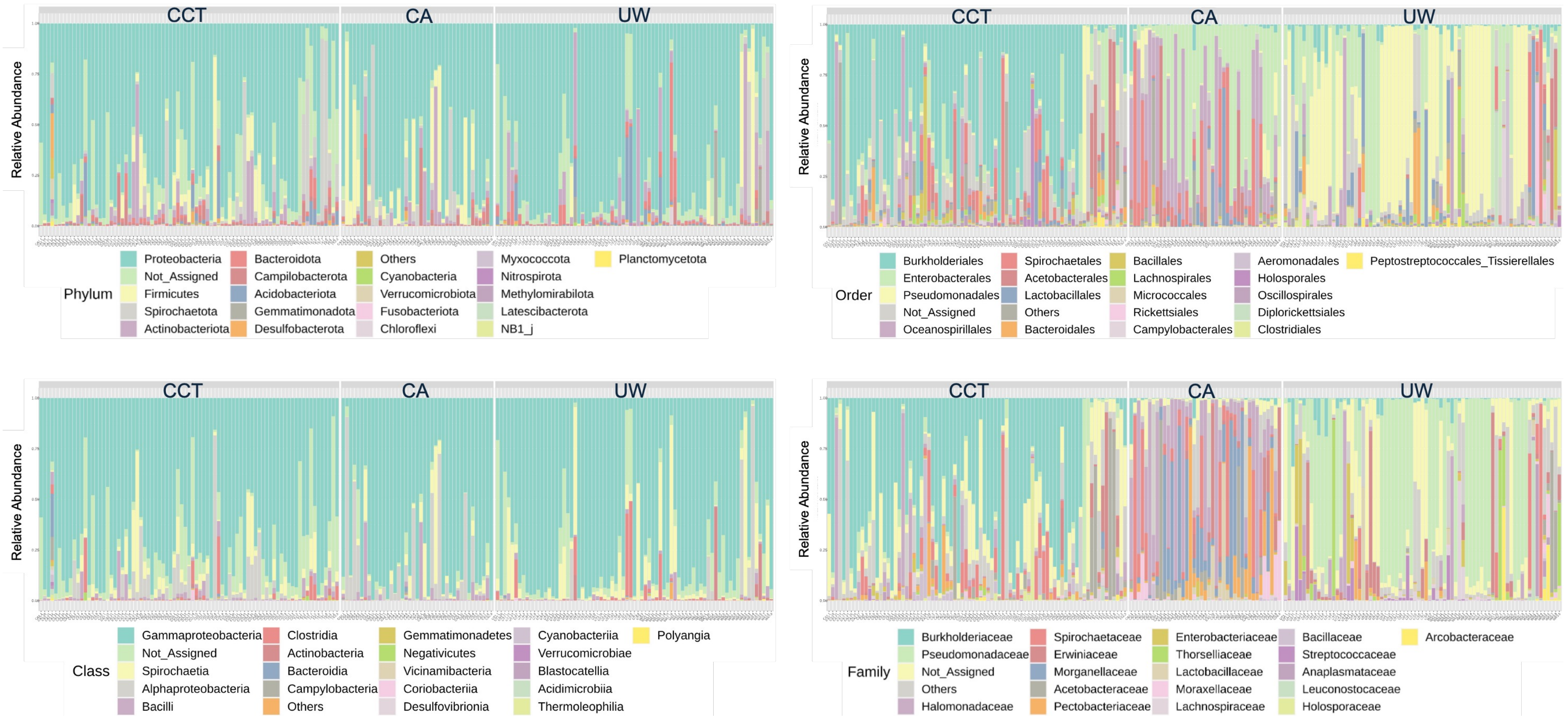


Additional file 3

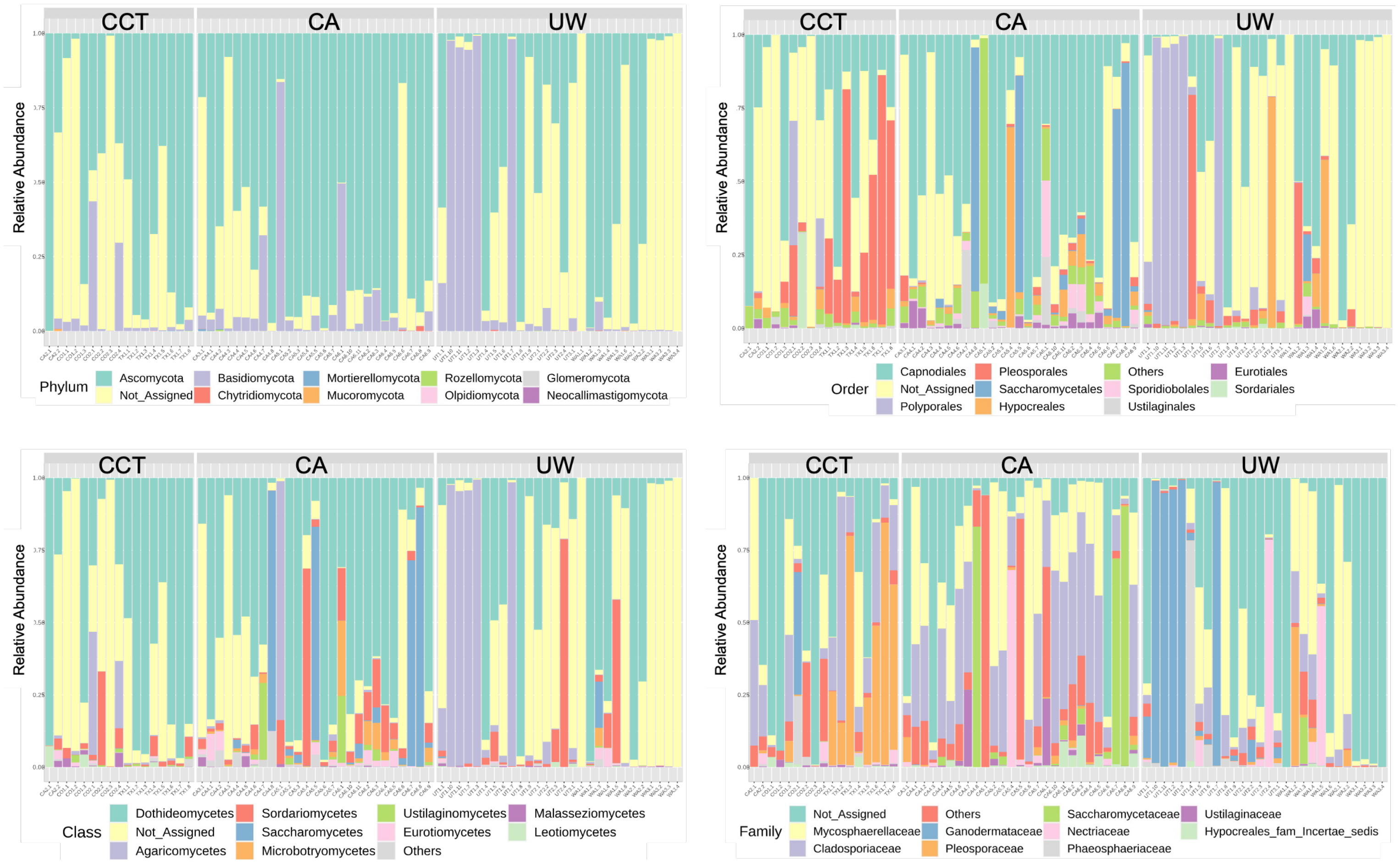


Additional file 4

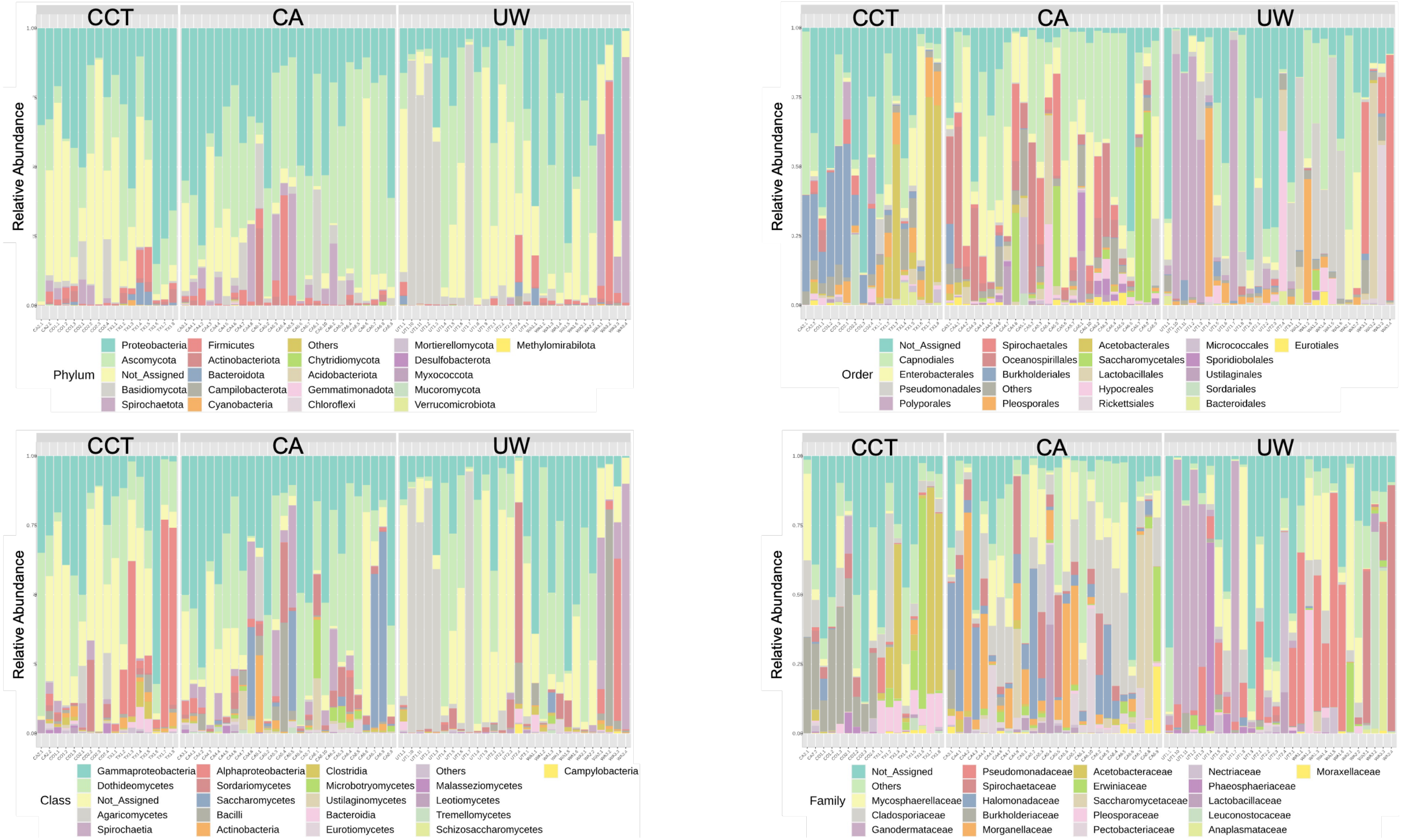


Additional file 5

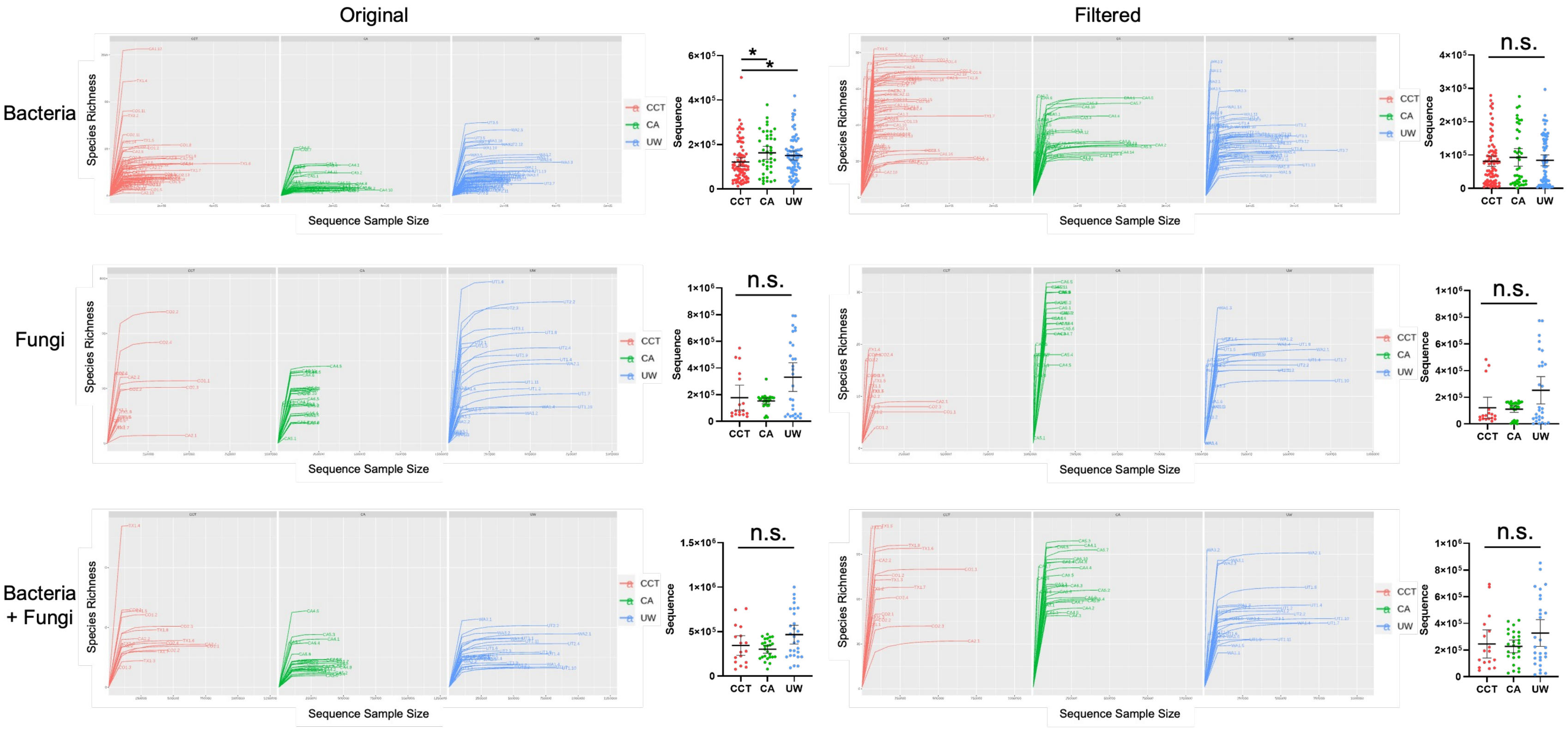


Additional file 6

**Bacteria**

The OOB error is 0.0863

| | CCT | CA | UW | class.error |
|---|---|---|---|---|
| CCT | 72 | 0 | 9.0 | 0.111 |
| CA | 1 | 39 | 1.0 | 0.0488 |
| UW | 6 | 0 | 69.0 | 0.08 |

The OOB error is 0.0914

| | CCT | CA | UW | class.error |
|---|---|---|---|---|
| CCT | 72 | 0 | 9.0 | 0.111 |
| CA | 1 | 39 | 1.0 | 0.0488 |
| UW | 7 | 0 | 68.0 | 0.0933 |

The OOB error is 0.0812

| | CCT | CA | UW | class.error |
|---|---|---|---|---|
| CCT | 72 | 0 | 9.0 | 0.111 |
| CA | 1 | 39 | 1.0 | 0.0488 |
| UW | 5 | 0 | 70.0 | 0.0667 |

The OOB error is 0.0863

| | CCT | CA | UW | class.error |
|---|---|---|---|---|
| CCT | 72 | 0 | 9.0 | 0.111 |
| CA | 1 | 39 | 1.0 | 0.0488 |
| UW | 6 | 0 | 69.0 | 0.08 |

The OOB error is 0.0863

| | CCT | CA | UW | class.error |
|---|---|---|---|---|
| CCT | 72 | 0 | 9.0 | 0.111 |
| CA | 1 | 39 | 1.0 | 0.0488 |
| UW | 6 | 0 | 69.0 | 0.08 |

**Fungi**

The OOB error is 0.222

| | CCT | CA | UW | class.error |
|---|---|---|---|---|
| CCT | 8 | 3 | 6.0 | 0.529 |
| CA | 3 | 24 | 0.0 | 0.111 |
| UW | 3 | 1 | 24.0 | 0.143 |

The OOB error is 0.222

| | CCT | CA | UW | class.error |
|---|---|---|---|---|
| CCT | 8 | 3 | 6.0 | 0.529 |
| CA | 3 | 24 | 0.0 | 0.111 |
| UW | 3 | 1 | 24.0 | 0.143 |

The OOB error is 0.222

| | CCT | CA | UW | class.error |
|---|---|---|---|---|
| CCT | 8 | 3 | 6.0 | 0.529 |
| CA | 3 | 24 | 0.0 | 0.111 |
| UW | 3 | 1 | 24.0 | 0.143 |

The OOB error is 0.236

| | CCT | CA | UW | class.error |
|---|---|---|---|---|
| CCT | 8 | 3 | 6.0 | 0.529 |
| CA | 3 | 24 | 0.0 | 0.111 |
| UW | 4 | 1 | 23.0 | 0.179 |

The OOB error is 0.236

| | CCT | CA | UW | class.error |
|---|---|---|---|---|
| CCT | 8 | 3 | 6.0 | 0.529 |
| CA | 3 | 24 | 0.0 | 0.111 |
| UW | 4 | 1 | 23.0 | 0.179 |

**Bacteria + Fungi**

The OOB error is 0.141

| | CCT | CA | UW | class.error |
|---|---|---|---|---|
| CCT | 13 | 0 | 4.0 | 0.235 |
| CA | 1 | 25 | 0.0 | 0.0385 |
| UW | 5 | 0 | 23.0 | 0.179 |

The OOB error is 0.155

| | CCT | CA | UW | class.error |
|---|---|---|---|---|
| CCT | 12 | 0 | 5.0 | 0.294 |
| CA | 1 | 25 | 0.0 | 0.0385 |
| UW | 5 | 0 | 23.0 | 0.179 |

The OOB error is 0.155

| | CCT | CA | UW | class.error |
|---|---|---|---|---|
| CCT | 12 | 0 | 5.0 | 0.294 |
| CA | 1 | 25 | 0.0 | 0.0385 |
| UW | 5 | 0 | 23.0 | 0.179 |

The OOB error is 0.155

| | CCT | CA | UW | class.error |
|---|---|---|---|---|
| CCT | 12 | 0 | 5.0 | 0.294 |
| CA | 1 | 25 | 0.0 | 0.0385 |
| UW | 5 | 0 | 23.0 | 0.179 |

The OOB error is 0.155

| | CCT | CA | UW | class.error |
|---|---|---|---|---|
| CCT | 12 | 0 | 5.0 | 0.294 |
| CA | 1 | 25 | 0.0 | 0.0385 |
| UW | 5 | 0 | 23.0 | 0.179 |

Additional file 7

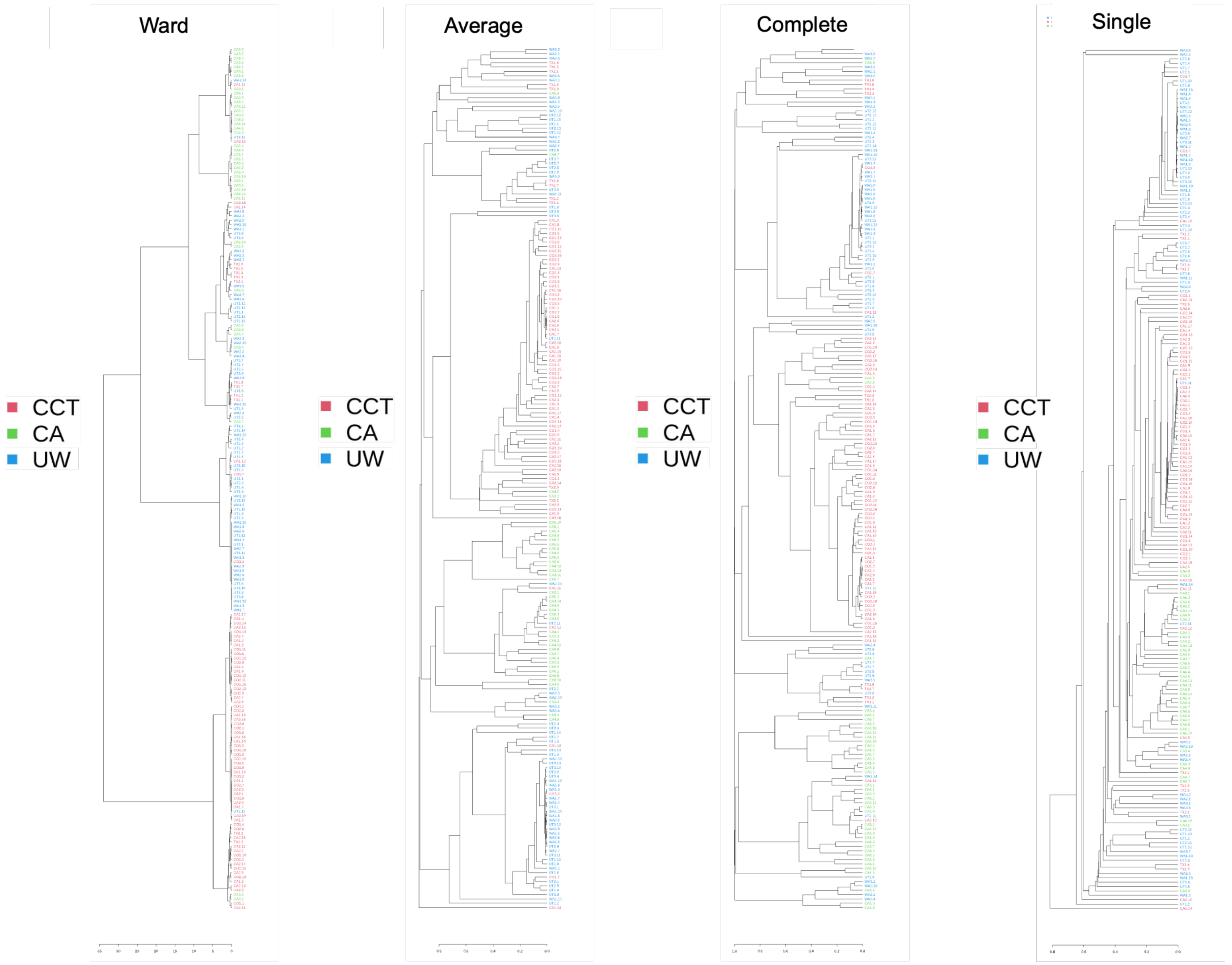


Additional file 8

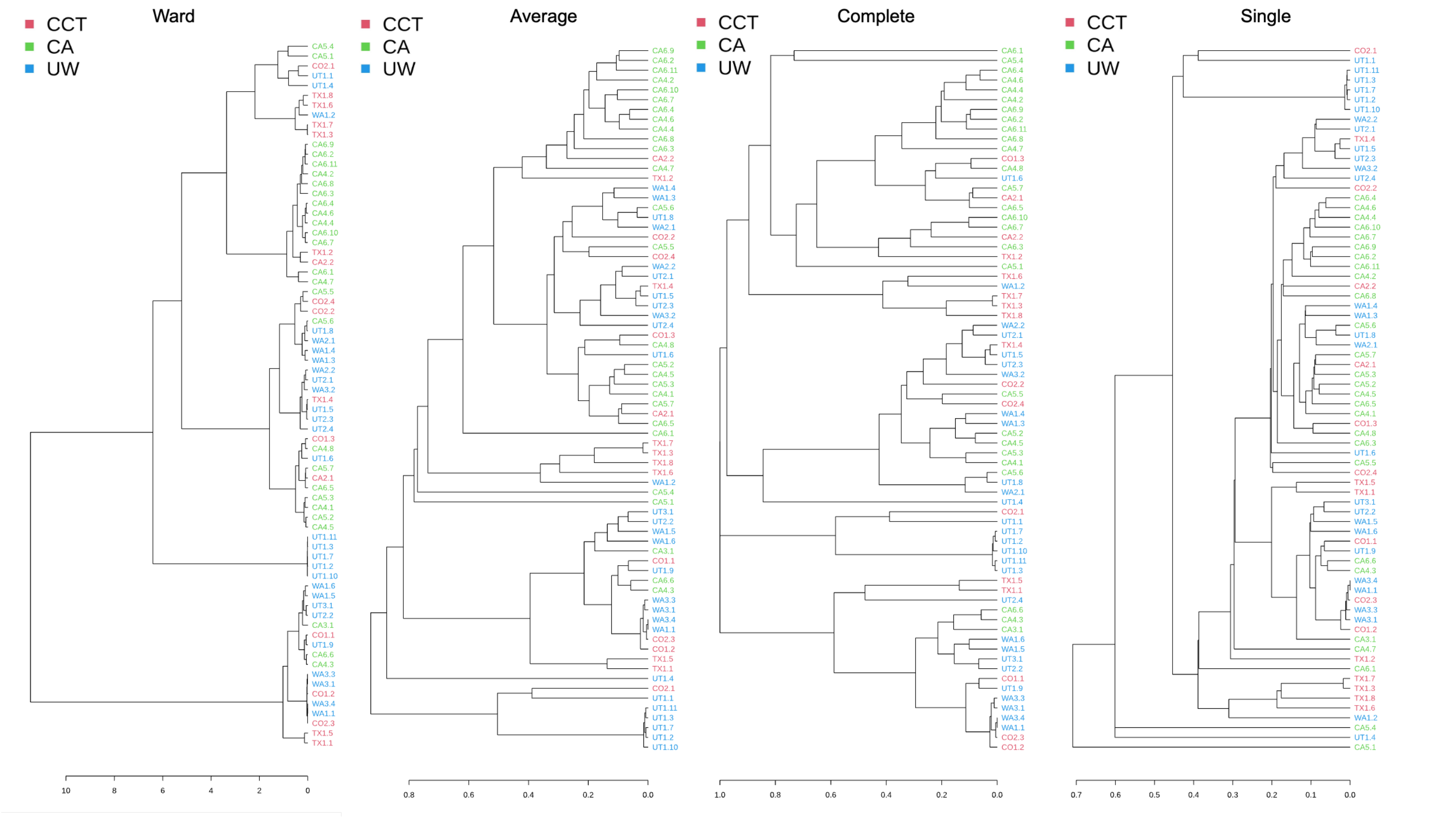


Additional file 9

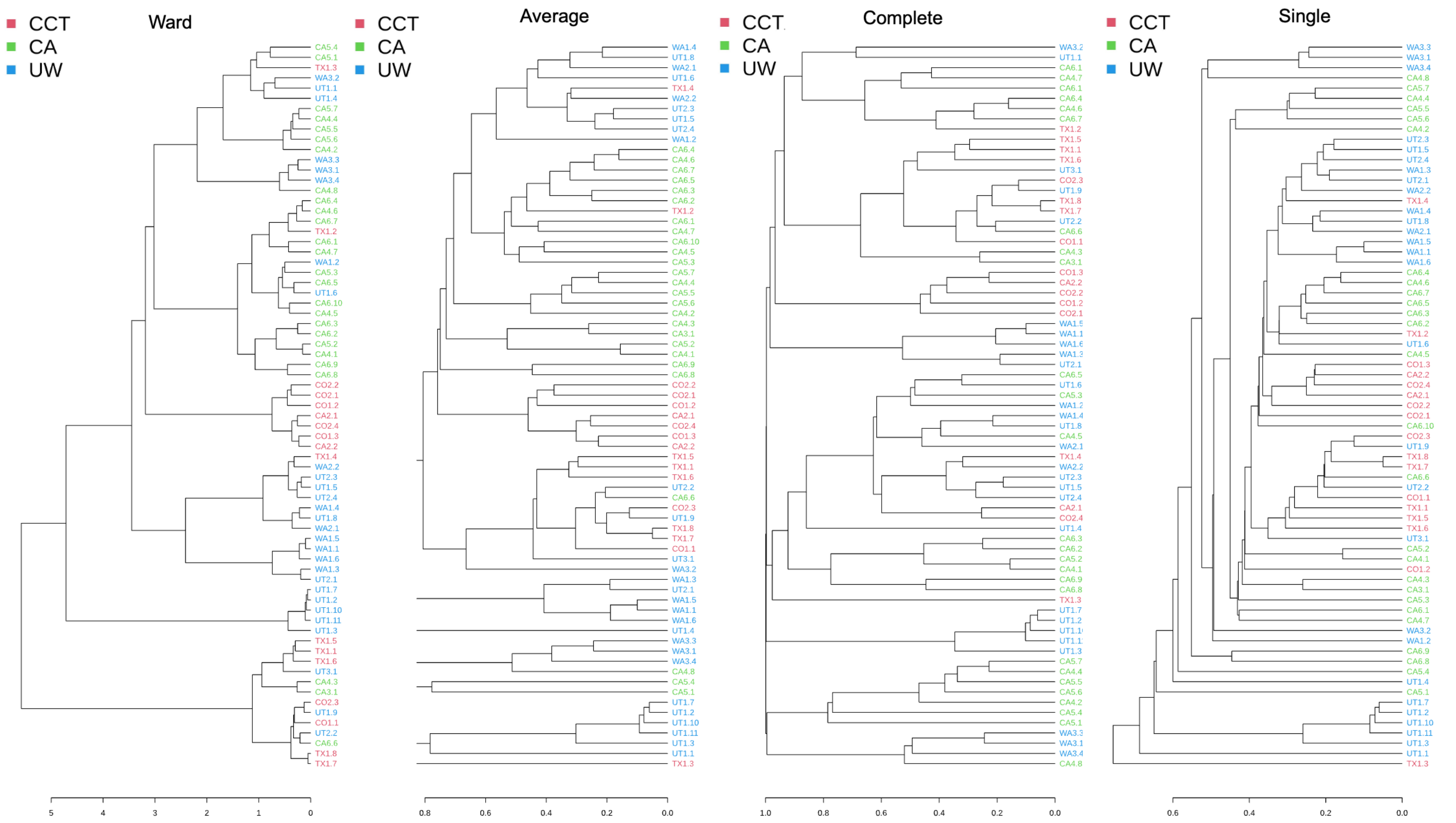


Additional file 10

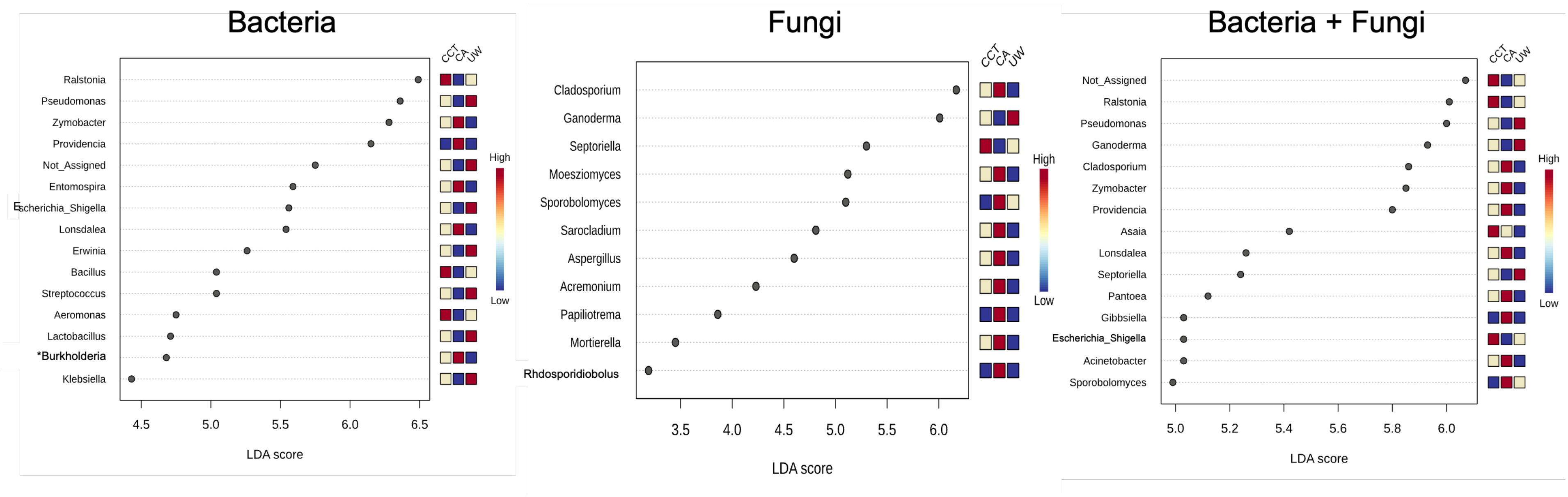


Additional file 11

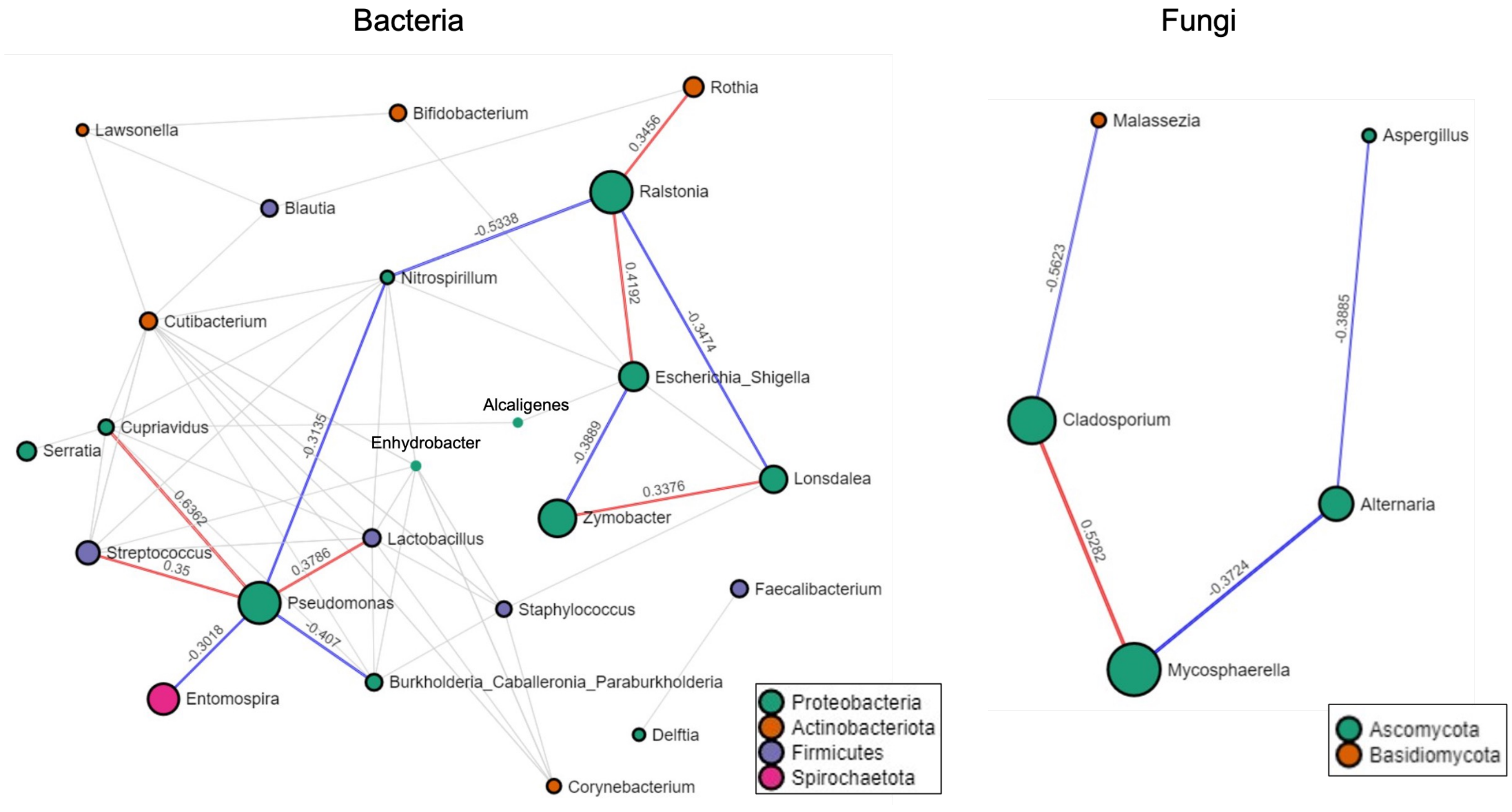


Additional file 12

**Additional File 13** Correlation network table for bacterial community

| Taxon1 | Taxon2 | Correlation | *P* value |
|---|---|---|---|
| *Alcaligenes* | *Escherichia_Shigella* | 0.3528 | 3.00E-04 |
| *Alcaligenes* | *Cupriavidus* | -0.3045 | 0.0049 |
| *Bifidobacterium* | *Escherichia_Shigella* | 0.3002 | 0.0017 |
| *Bifidobacterium* | *Lawsonella* | -0.3734 | 0.0039 |
| *Blautia* | *Cutibacterium* | -0.3215 | 5.00E-04 |
| *Blautia* | *Rothia* | -0.3276 | 0.0053 |
| *Blautia* | *Lawsonella* | -0.3398 | 0.0056 |
| *Burkholderia_Caballeronia_Paraburkholderia* | *Cutibacterium* | 0.3264 | 0 |
| *Burkholderia_Caballeronia_Paraburkholderia* | *Enhydrobacter* | 0.4071 | 0 |
| *Burkholderia_Caballeronia_Paraburkholderia* | *Pseudomonas* | -0.407 | 0 |
| *Burkholderia_Caballeronia_Paraburkholderia* | *Staphylococcus* | 0.3472 | 1.00E-04 |
| *Burkholderia_Caballeronia_Paraburkholderia* | *Cupriavidus* | 0.3416 | 2.00E-04 |
| *Burkholderia_Caballeronia_Paraburkholderia* | *Lactobacillus* | -0.3346 | 5.00E-04 |
| *Corynebacterium* | *Cutibacterium* | 0.4959 | 0 |
| *Corynebacterium* | *Enhydrobacter* | 0.6077 | 0 |
| *Corynebacterium* | *Lactobacillus* | -0.4107 | 0 |
| *Corynebacterium* | *Staphylococcus* | 0.431 | 0 |
| *Cupriavidus* | *Cutibacterium* | -0.4146 | 0 |
| *Cupriavidus* | *Not_Assigned* | -0.369 | 0 |
| *Cupriavidus* | *Pseudomonas* | 0.6362 | 0 |
| *Cupriavidus* | *Streptococcus* | 0.4627 | 0 |
| *Cupriavidus* | *Lactobacillus* | 0.3716 | 1.00E-04 |
| *Cupriavidus* | *Burkholderia_Caballeronia_Paraburkholderia* | 0.3416 | 2.00E-04 |
| *Cupriavidus* | *Serratia* | -0.3417 | 3.00E-04 |
| *Cupriavidus* | *Nitrospirillum* | 0.3373 | 4.00E-04 |
| *Cupriavidus* | *Alcaligenes* | -0.3045 | 0.0049 |
| *Cutibacterium* | *Burkholderia_Caballeronia_Paraburkholderia* | 0.3264 | 0 |
| *Cutibacterium* | *Corynebacterium* | 0.4959 | 0 |
| *Cutibacterium* | *Cupriavidus* | -0.4146 | 0 |
| *Cutibacterium* | *Enhydrobacter* | 0.555 | 0 |
| *Cutibacterium* | *Lactobacillus* | -0.4538 | 0 |
| *Cutibacterium* | *Lawsonella* | 0.4456 | 0 |
| *Cutibacterium* | *Nitrospirillum* | 0.3506 | 0 |
| *Cutibacterium* | *Staphylococcus* | 0.5704 | 0 |
| *Cutibacterium* | *Streptococcus* | -0.5522 | 0 |
| *Cutibacterium* | *Blautia* | -0.3215 | 5.00E-04 |
| *Delftia* | *Faecalibacterium* | 0.3266 | 0.0012 |
| *Enhydrobacter* | *Burkholderia_Caballeronia_Paraburkholderia* | 0.4071 | 0 |
| *Enhydrobacter* | *Corynebacterium* | 0.6077 | 0 |
| *Enhydrobacter* | *Cutibacterium* | 0.555 | 0 |
| *Enhydrobacter* | *Lactobacillus* | -0.4325 | 0 |
| *Enhydrobacter* | *Nitrospirillum* | 0.4525 | 0 |
| *Enhydrobacter* | *Staphylococcus* | 0.4983 | 0 |
| *Enhydrobacter* | *Streptococcus* | -0.3233 | 0.0019 |
| *Entomospira* | *Pseudomonas* | -0.3018 | 0.0019 |
| *Escherichia_Shigella* | *Lonsdalea* | -0.4638 | 0 |
| *Escherichia_Shigella* | *Nitrospirillum* | -0.3568 | 0 |
| *Escherichia_Shigella* | *Ralstonia* | 0.4192 | 0 |
| *Escherichia_Shigella* | *Zymobacter* | -0.3889 | 0 |
| *Escherichia_Shigella* | *Alcaligenes* | 0.3528 | 3.00E-04 |
| *Escherichia_Shigella* | *Bifidobacterium* | 0.3002 | 0.0017 |
| *Faecalibacterium* | *Delftia* | 0.3266 | 0.0012 |
| *Lactobacillus* | *Corynebacterium* | -0.4107 | 0 |
| *Lactobacillus* | *Cutibacterium* | -0.4538 | 0 |
| *Lactobacillus* | *Enhydrobacter* | -0.4325 | 0 |
| *Lactobacillus* | *Nitrospirillum* | -0.4426 | 0 |
| *Lactobacillus* | *Pseudomonas* | 0.3786 | 0 |
| *Lactobacillus* | *Streptococcus* | 0.4823 | 0 |
| *Lactobacillus* | *Cupriavidus* | 0.3716 | 1.00E-04 |
| *Lactobacillus* | *Staphylococcus* | -0.3852 | 2.00E-04 |
| *Lactobacillus* | *Burkholderia_Caballeronia_Paraburkholderia* | -0.3346 | 5.00E-04 |
| *Lawsonella* | *Cutibacterium* | 0.4456 | 0 |
| *Lawsonella* | *Bifidobacterium* | -0.3734 | 0.0039 |
| *Lawsonella* | *Blautia* | -0.3398 | 0.0056 |
| *Lonsdalea* | *Escherichia_Shigella* | -0.4638 | 0 |
| *Lonsdalea* | *Ralstonia* | -0.3474 | 7.00E-04 |
| *Lonsdalea* | *Zymobacter* | 0.3376 | 7.00E-04 |
| *Lonsdalea* | *Not_Assigned* | -0.3344 | 0.0015 |
| *Lonsdalea* | *Staphylococcus* | -0.341 | 0.0017 |
| *Nitrospirillum* | *Cutibacterium* | 0.3506 | 0 |

| | | | |
|---|---|---|---|
| *Nitrospirillum* | *Enhydrobacter* | 0.4525 | 0 |
| *Nitrospirillum* | *Escherichia_Shigella* | -0.3568 | 0 |
| *Nitrospirillum* | *Lactobacillus* | -0.4426 | 0 |
| *Nitrospirillum* | *Ralstonia* | -0.5338 | 0 |
| *Nitrospirillum* | *Streptococcus* | -0.4242 | 0 |
| *Nitrospirillum* | *Pseudomonas* | -0.3135 | 2.00E-04 |
| *Nitrospirillum* | *Cupriavidus* | 0.3373 | 4.00E-04 |
| *Not_Assigned* | *Cupriavidus* | -0.369 | 0 |
| *Not_Assigned* | *Streptococcus* | -0.3949 | 0 |
| *Not_Assigned* | *Lonsdalea* | -0.3344 | 0.0015 |
| *Pseudomonas* | *Burkholderia_Caballeronia_Paraburkholderia* | -0.407 | 0 |
| *Pseudomonas* | *Cupriavidus* | 0.6362 | 0 |
| *Pseudomonas* | *Lactobacillus* | 0.3786 | 0 |
| *Pseudomonas* | *Streptococcus* | 0.35 | 0 |
| *Pseudomonas* | *Nitrospirillum* | -0.3135 | 2.00E-04 |
| *Pseudomonas* | *Entomospira* | -0.3018 | 0.0019 |
| *Ralstonia* | *Escherichia_Shigella* | 0.4192 | 0 |
| *Ralstonia* | *Nitrospirillum* | -0.5338 | 0 |
| *Ralstonia* | *Rothia* | 0.3456 | 4.00E-04 |
| *Ralstonia* | *Lonsdalea* | -0.3474 | 7.00E-04 |
| *Rothia* | *Ralstonia* | 0.3456 | 4.00E-04 |
| *Rothia* | *Blautia* | -0.3276 | 0.0053 |
| *Serratia* | *Cupriavidus* | -0.3417 | 3.00E-04 |
| *Staphylococcus* | *Corynebacterium* | 0.431 | 0 |
| *Staphylococcus* | *Cutibacterium* | 0.5704 | 0 |
| *Staphylococcus* | *Enhydrobacter* | 0.4983 | 0 |
| *Staphylococcus* | *Burkholderia_Caballeronia_Paraburkholderia* | 0.3472 | 1.00E-04 |
| *Staphylococcus* | *Lactobacillus* | -0.3852 | 2.00E-04 |
| *Staphylococcus* | *Lonsdalea* | -0.341 | 0.0017 |
| *Streptococcus* | *Cupriavidus* | 0.4627 | 0 |
| *Streptococcus* | *Cutibacterium* | -0.5522 | 0 |
| *Streptococcus* | *Lactobacillus* | 0.4823 | 0 |
| *Streptococcus* | *Nitrospirillum* | -0.4242 | 0 |
| *Streptococcus* | *Not_Assigned* | -0.3949 | 0 |
| *Streptococcus* | *Pseudomonas* | 0.35 | 0 |
| *Streptococcus* | *Enhydrobacter* | -0.3233 | 0.0019 |
| *Zymobacter* | *Escherichia_Shigella* | -0.3889 | 0 |
| *Zymobacter* | *Lonsdalea* | 0.3376 | 7.00E-04 |

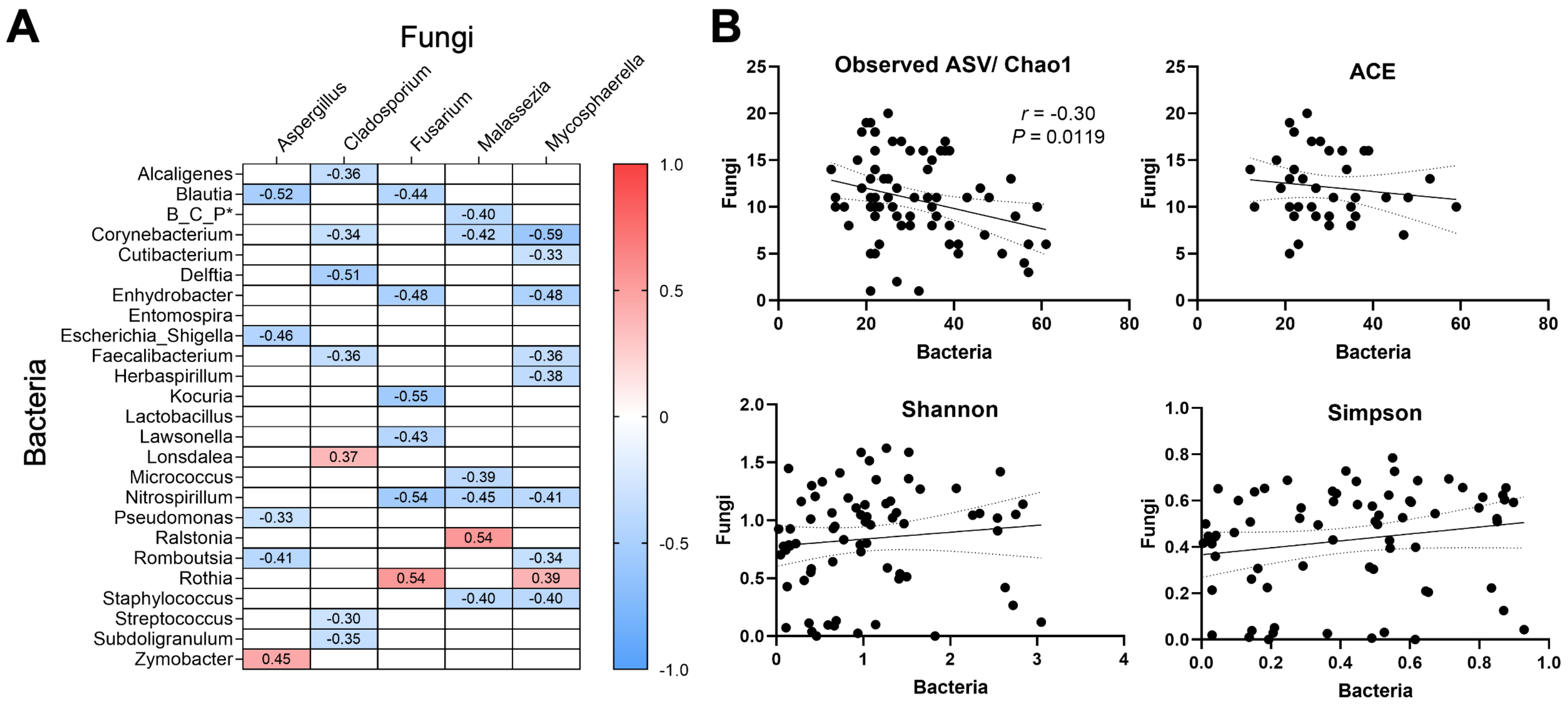


Additional file 14
